# A Conditional Variational Autoencoder with QSAR-Guided Surrogate-Weighted Fine-Tuning and Cross-Entropy Optimization for Targeted Antimicrobial Peptide Generation

**DOI:** 10.64898/2026.04.28.721185

**Authors:** Ismael Castanon, Fangping Wan, César de la Fuente-Núñez, Alessandro Pini, Chiara Falciani

**Affiliations:** Department of Chemistry, Bioscience and Environmental Engineering, University of Stavanger, Stavanger, Norway; Machine Biology Group, Departments of Psychiatry and Microbiology, Institute for Biomedical Informatics, Institute for Translational Medicine and Therapeutics, Perelman School of Medicine, University of Pennsylvania, Philadelphia, Pennsylvania, United States of America; Departments of Bioengineering and Chemical and Biomolecular Engineering, School of Engineering and Applied Science, University of Pennsylvania, Philadelphia, Pennsylvania, United States of America; Department of Chemistry, School of Arts and Sciences, University of Pennsylvania, Philadelphia, Pennsylvania, United States of America; Penn Institute for Computational Science, University of Pennsylvania, Philadelphia, Pennsylvania, United States of America; SetLance, SRL, Via Fiorentina 1, 53100 Siena, Italy; Department of Medical Biotechnology, University of Siena, Italy; Clinical Pathology Unit, Azienda Ospedaliera, Universitaria Senese, Siena

## Abstract

Machine Learning frameworks have emerged as a promising tool for antimicrobial peptide design; however, generative models remain limited by two persistent problems: the limited availability of experimentally validated peptides and the circular dependency of the models. In this work we present a conditional variational autoencoder pipeline that addresses both limitations through a modular architecture that combines both binary and quantitative experimental data and implements a multimodal approach to externally guide the generation. A transformer-based encoder successfully generated a discriminative 64-dimensional latent space (test AUROC 0.968, F1 0.919) separating antimicrobial from non-antimicrobial sequences. This latent representation conditions a species-specific LoRA fine-tuned ProtGPT2 decoder through a scalar gating function, which generates balanced antimicrobial peptides through two different modes; prior and perturb, depending on their generation starting points. We introduced a Surrogate Weighted Fine-Tuning (SWF) ensemble to eliminate the circular dependency and a Cross-Entropy Method to explore and exploit the latent space, leading to successful antimicrobial peptide generation. The best candidates exhibited competitive physicochemical characteristics, a mean helical fraction of 0.874 (mean pLDDT 83.7), and externally predicted efficacy evaluated by APEX.

## Introduction

Antimicrobial peptides (AMPs) have emerged as a promising alternative to conventional antibiotics in the context of the global antimicrobial resistance (AMR) crisis.^1,2^ Antimicrobial peptides are present in several organisms, from bacteria or fungus to humans, and possess an action mechanism that primarily targets the bacterial wall.^3^ Different action mechanisms such as barrel-stave, toroidal, and carpet model have been proposed, but all of them necessitate similar physicochemical characteristics such as helicity, hydrophobicity and a high cationic charge.^4,5^ Despite their potential, clinical translation has exhibited several problems, due to their poor pharmacology properties such as low blood stability, bad distribution rates, and high cytotoxicity.^6–9^ Numerous strategies have been proposed to address these bottlenecks, including the incorporation of D-amino acids, peptide cyclization, and conjugation with hydrophobic moieties such as polyethylene glycol, to name some of them.^10–13^ While effective, these strategies addressed pharmacological bottlenecks rather than optimizing the sequence itself. Classical rational design based on physicochemical descriptors and empirical evidence has provided useful mechanistic insight but explores only a negligible fraction of the combinatorial sequence space available to peptide designers.

During the last years, artificial intelligence, and specifically neural networks have emerged as a powerful tool for navigating this space. Generative models including variational autoencoders (VAEs)^14^, generative adversarial networks (GANs)^15^, and Large Language Models (LLMs)^16,17^ have demonstrated the ability to produce novel AMP sequences with predicted antimicrobial activity. However, existing approaches face two persistent limitations. First, experimentally validated MIC data remain scarce, forcing most models to rely on binary active/inactive labels rather than continuous activity values, which limits their ability to optimize toward specific potency targets. Second, and more fundamentally, most generative pipelines use the same model or closely related features for both generation and evaluation, creating a circular dependency that inflates apparent performance without guaranteeing improvement in independently scored candidates.

Here we present a conditional variational autoencoder (CVAE) pipeline that addresses both limitations through a modular optimization strategy. A transformer-based encoder is first trained to generate a discriminative latent space separating antimicrobial and non-antimicrobial peptide sequences. This latent representation conditions a ProtGPT2 decoder species-specific fine-tuned via Low-Rank Adaptation (LoRA)^18^ with a learned prefix conditioning mechanism. Pareto Fronts were built to evaluate optimal generation regimes through two different generation modes; prior and perturb. Later in the pipeline, we implemented a Surrogate-Weighted Fine-Tuning (SWF) strategy based on an external QSAR-based ensemble trained independently on experimental MIC data, eliminating the circular dependency present in prior work.^19,20^ We further implemented Cross-Entropy Method (CEM) optimization over the latent space proving it as a useful strategy depending on the latent geometry, a finding with direct implications for the design of iterative generative pipelines^21^.

## Methods

### Encoder Architecture and Training

The encoder was implemented as a Transformer-based variational autoencoder trained to organize the latent space to discriminate antimicrobial from non-antimicrobial peptide sequences. The architecture consists of four self-attention layers containing eight heads each, a model dimensionality of 256 dimensions converted into a 64-dimensional latent space by projecting CLS token representation through a reparameterization bottleneck.^22,23^ The classification head was set to operate on the sample vector z instead of the posterior mean μ. For downstream conditioning in the CVAE and for the subsequent latent-space analyses the deterministic posterior mean μ was used as the encoder representation. A free-bits KL penalty was implemented to avoid posterior collapse, and word dropout was also present to avoid overfitting.^24^ The model was evaluated for 20 epochs implementing BCE-based early stopping, to prioritize reconstruction. The dataset was built from scratch utilizing data from DBAASP, DRAMP, APD3 and CAMPR4. It comprises 5,000 peptide sequences with binary AMP/noAMP labels, exhibiting balanced positive and negative sets (≈49% AMP across all splits). To prevent homology-based data leakage, sequences were clustered using MMseqs2^25^ with a minimum sequence identity threshold of 0.70 and a coverage threshold of 0.80. Cluster assignments were used to partition the dataset at the cluster level rather than the sequence level, ensuring that no homologous sequences appeared across training, validation, and test splits. This yielded 3,483 training sequences (1,696 AMP / 1,787 noAMP), 776 validation sequences (394 AMP / 382 noAMP), and 741 test sequences (358 AMP / 383 noAMP).

### CVAE Architecture and Training

ProtGPT2 was used as decoder, it is a GPT-2-based transformer language model comprising 36 layers and approximately 738 million parameters, pre-trained on approximately 50 million non-annotated protein sequences from the UniRef50 database using a causal language modeling objective.^26^ At inference time, a latent vector z is passed through the prefix MLP and the resulting prefix embeddings are prepended to the decoder input, together with the bacterial condition token. The decoder then generates sequences autoregressively, sampling from the output distribution at each position according to a temperature parameter T that controls the sharpness of the sampling distribution. The complete Conditional Variational AutoEncoder (CVAE) was completed by connecting the encoder with a ProtGPT2 decoder fine-tuned implementing a combination of a Low-Rank Adaptation (LoRA) and a prefix conditioning mechanism.^18^ The latent space information flowed into the decoder regularized by a scalar function which projected a learned prefix of length 16 projected from μ via a two-layer MLP, prepended to the decoder input embeddings and processed through all attention layers as soft prompt tokens.^27^ A floor value of 0.2 was set to prevent complete prefix shutdown during training. The gate was computed from mean μ to ensure deterministic conditioning instead of utilizing the sampled z.

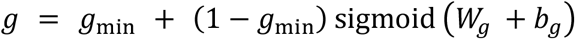

Four free-bits regimes were evaluated (0.01, 0.03, 0.10, and 0.30) to assess the most informative schedule.^24^ The loss function computed a language modeling term, a regression term penalizing deviation from target activity scores, a classification term, and the KL divergence, weighted by a cyclical beta schedule that annealed from 0 to 1 over training. The model selection was based on a composite validation score that balanced language modelling, regression and classification losses. This score composite was set up based on previous analyses (not shown) and does not include the number of dimensions to influence in the selected epoch, since the free-bits and the gate dynamics already proved the informative capacity of the latent. The regression and classification heads were attached to the decoder final hidden state.

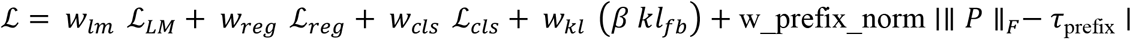

The dataset was built from scratch utilizing data from DBAASP, DRAMP, APD3 and CAMPR4. It comprises 2,829 unique peptide sequences with experimentally derived MIC values expressed as log-transformed µg/mL. It comprises a common set of negative peptides against all the three bacteria (MIC 4.0) of 1512 sequences, built as commented before in this work. The positive species-specific set comprises 624 peptides against *E. coli*, 602 peptides against *P. aeruginosa*, and 261 peptides against *K. pneumoniae*.

### Base CVAE Generation

The generation from the decoder was evaluated across a grid of sampling modes including perturbation magnitudes and decoding temperatures through the construction of Pareto Fronts^28^ to compare the two generation regimes implemented; prior, in which latent vectors are drawn directly from the standard Gaussian prior, and perturb, in which a selected seed is coded by the encoder utilizing mean μ as a starting point, and subsequently displace Gaussian noise of magnitude σ before decoding. For reproducibility and fair comparison, we utilized the experimentally tested L33 sequence (KKIRVRLSA) as a seed all through this word. The Pareto fronts were constructed utilizing σ values of 0.05, 0.20, and 0.40 for perturb mode, 0.30, 0.50, and 0.70 for prior mode, and decoding temperatures of 0.75, 0.85, and 0.95, under each of the three bacterial conditions.

Each regime was evaluated through two independent random seeds, yielding 36 distinct regime combinations and a total of 5,400 generated antimicrobial sequences. Those sequences were then scored post hoc to calculate their OOD, hit rate and diversity. The Out-of-Distribution metric was defined by hard-constraints such as sequence length (8–25 residues), number of cationic residues (2–10), and GRAVY index (−1.5 to 2.5).^29^ The diversity was computed calculating mean 3-mer novelty relative to the training set, and the hit rate was computed by the QSAR-based ensemble utilizing the external score (hit rate at predicted MIC ≤ 1 μg/mL). The regime selection was formulated as a multi-objective optimization problem containing these three mentioned competing objectives to maximize diversity and hit-rate, while minimizing the out-of-distribution risk. Both generation modes prior and perturb were evaluated separately.

From the 18 unique regime combinations evaluated across both modes, 11 of them shaped the pareto front, corresponding to 61% of the explored landscape. Within the perturb mode, 7 out of 9 configurations were non-dominated, while only 4 out of 9 were non-dominated for the prior generation mode. The selected perturb regime for downstream experiments (σ = 0.20, T = 0.95), was extracted from this Pareto Front as the primary generation configuration for fine tuning and factorial evaluation, together with two different prior regimes (T = 0.95) differing in their sigma value (σ = 0.3, 0.70).

### QSAR-Guided Surrogate-Weighted Fine-Tuning (SWF)

The QSAR ensemble was trained on experimentally derived MIC data to provide a fixed scoring function independent of the generative model. The ensemble consisted of five multilayer perceptron models; each trained on 25 physicochemical descriptors selected by minimum Redundancy Maximum Relevance (mRMR) from a pool of 50 sequence-derived features computed using the Peptidy library at pH 7.0.^30,31^ To include the uncertainty component a conservative upper-confidence bound score was computed for activity predictions as the ensemble mean plus one standard deviation, therefore penalizing uncertain predictions and biases selection towards high-confidence candidates.

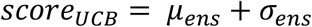

The fine-tuning dataset was assembled from 1,716 baseline-generated candidates scored by the QSAR ensemble.^19,20^ A two-stage filtering procedure was applied: first, hard physicochemical constraints removed sequences outside biologically plausible ranges for length (8–25 residues), cationic residue ratio (0.15–0.55), net charge (0–12), and GRAVY index (−2.5 to 1.0), retaining 1,591 sequences. Second, candidates were ranked by a penalized score defined as the conservative QSAR upper-confidence-bound score plus soft penalties for deviation from target length, cationic ratio, and GRAVY, and the top 30% under this penalized score were retained, yielding 478 sequences of which 470 were unique. Per-sequence training weights were then computed from the same penalized score and rescaled to the interval [0.1, 1], assigning higher weight to candidates with lower penalized scores.

The base CVAE was fine-tuned on this weighted dataset for a maximum of 120 epochs with early stopping based on validation loss. The best checkpoint was obtained at epoch 6 with a validation loss of 3.464. To ensure a fair comparison, all model outputs were rescored using the same fixed external QSAR ensemble.

### Latent Space Optimization via CEM and Comparative Branches

Cross-Entropy Method (CEM) optimization was applied over the 64-dimensional latent space.^21^ At each iteration, a population of latent vectors was sampled from a Gaussian distribution centered at the current mean μ with per-dimension standard deviation σ. The generated sequences were decoded, and scored by the external ensemble, which penalizes uncertain predictions and favors high-confidence candidates. The top-scoring elite fraction candidates are then retained, for the subsequent CEM cycle. The optimizer was run for 20 iterations with a population of 200 sequences each, with a fixed number of 40 sequences per iteration, retained in a buffer of up to 500 sequences across iterations. The distribution mean was initialized at μ = 0 updated with a smoothing factor of 0.5 while following an adaptive schedule aiming to sample variance (σ from 0.95 to 1.20) and gate scale (0.82 to 1.05) across iterations. All comparisons were performed under a fixed external scoring protocol evaluating the top 250 candidates per pipeline branch.

A REINFORCE-based reinforcement learning framework was also evaluated utilizing the external score as the scalar reward.^32^ At every training step, the initialized batch of elite vectors sampled through CEM was then used for generation. The scalar reward was defined as the negative conservative external score, plus a weighting term based on physicochemical plausibility, minus a repetition penalty proportional to the maximum 3-mer count in the generated sequence. Advantages were computed relative to a running exponential baseline exhibiting a 0.95 decay, while the policy loss was augmented with KL penalty against the frozen reference model in order to prevent collapse (weighted by a coefficient of 0.2). The LoRA adapter weights were optimized using AdamW with a learning rate of 2×10^−6^ and gradient clipping at unit norm for 100 steps with a batch size of 8.

### Candidate Characterization

The best selected model: CEM generation decoding after reweighting through external scorer (trained on perturb candidates), was used to generate 500 sequences against *E. coli*, from which 465 were unique. From those sequences, the top 100 exhibiting the best external score were selected for subsequent analyses. To contextualize the physicochemical profile of the synthetic antimicrobial peptides, we first extracted a Gram-negative antimicrobial dataset from DBAASP composed by non-redundant antimicrobial sequences exhibiting a length between 8 to 30 natural amino acids, yielding 1,336 peptides. Several physicochemical characteristics were computed using modlamp 4.3.2. Net charge was calculated at pH 7.4 using the Henderson-Hasselbalch formulation. The grand average of hydrophobicity (GRAVY) was computed using the Kyte-Doolittle scale. The hydrophobic moment was calculated using the Eisenberg consensus scale with a helical wheel angle of 100 degrees across the full sequence length. The cationic residue ratio was defined as the fraction of lysine and arginine residues relative to total sequence length.

ESMFold v1 was used to predict the three-dimensional structures of the candidates, so we could also analyze parameters such as the helical content through Biopython PPBuilder module.^33^ A residue was classified as alpha-helical if its φ angle fell within [−80°, −40°] and its ψ angle within [−60°, −20°], corresponding to the alpha-helical region of the Ramachandran diagram. Helicity was expressed as the fraction of residues with valid φ/ψ assignments satisfying this criterion.

## Results

### Encoder Performance

Classification performance on the validation set remained stable across training, with AUROC ranging between 0.92 and 0.95 and AUPRC between 0.93 and 0.955. Epoch 9 was selected as primary checkpoint corresponding to an AUROC validation of 0.949. The test set exhibited an AUROC of 0.968, AUPRC of 0.961, accuracy of 0.924, and F1 of 0.919, confirming that performance generalized well beyond the validation split.

The KL diagnostic per dimension increased gradually from approximately 0.50 at epoch zero to around 0.65-0.75 forming a plateau that showed a final test value of 0.753. The effective dimensionality at the selected checkpoint was 61.1 out of an existing 64 latent space dimensions. The mean latent standard deviation of 0.575 illustrates that posterior was healthy, showing no significant deviation, consistent with a well-regularized representation.

The PCA projection of the latent means revealed a discriminative geometric structure in which AMP and non-AMP sequences occupied clearly distinct regions of the two principal components. Non-antimicrobial form a compact, dense cluster at high positive values of the first principal component while antimicrobial sequences are distributed at negative values. The distribution of latent norms ‖μ‖ also provided valuable insights about antimicrobial latent geometry, concentrating non-AMP sequences in a unimodal distribution centered around 6-7, while AMP sequences densely concentrated at the upper norm boundary close to ten. This structural asymmetry suggests that antimicrobial activity is encoded as high-norm, boundary-proximal vectors, which would determine the subsequent generation pipeline.

**Figure 1.**
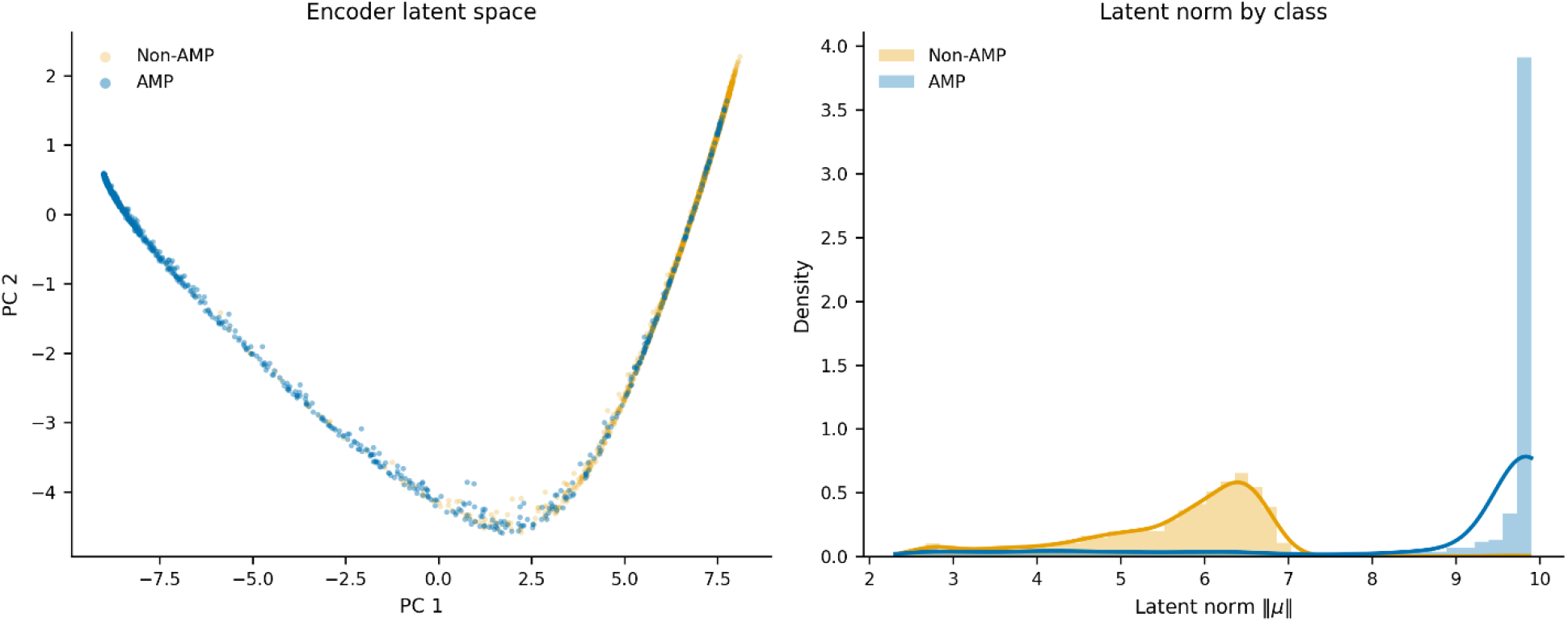
PCA latent representation and latent norm by class. Principal-component projection of the latent space exhibited a discriminative gradient between antimicrobial and non-antimicrobial sequences. The latent-norm distributions further revealed an asymmetric geometry, concentrating antimicrobial candidates in boundary-proximal regions of the latent space.

### CVAE Training Dynamics and Gate Behavior

As previously commented in the methods section, the composite score combining language degradation, classification and regression heads determined the selected checkpoint, reaching its minimum at epoch 9, after which all component losses began to degrade. The language modeling component, which was initialized from ProtGPT2 showed slight improvement during the first 5 epochs, followed by a progressive deterioration. Regression and classification heads showed an initial rapid improvement, leading into a plateau around epochs 6-10, followed by a deterioration thereafter. Four different free bits regimes were evaluated to get a better picture of the information flow through the prefix. The free bits regime 0.30 achieved the best raw language modeling loss (val_lm = 5.39) but forced a KL of 11.57 nats and compressed the gate distribution artificially, with gate standard deviation collapsing to 0.059. At the opposite end, the lower free bits regime 0.01 showed the worst language modelling while presenting a KL of 0.35 nats, exhibiting an almost inert prefix channel, as it can be observed at the direct conditioning figure, which shows practically 0 difference through the condition and shuffle ablation experiments. The free bits regime 0.1 represented a moderate pathway between both extremes, exhibiting the best prefix off ablation experiment values, a KL of 4.35 nats, and a gate standard deviation of 0.209. These results support the idea that 0.1 free bits scheme could be close to the optimal size to maintain the compromise between regularization and informative potential.

**Figure 2.**
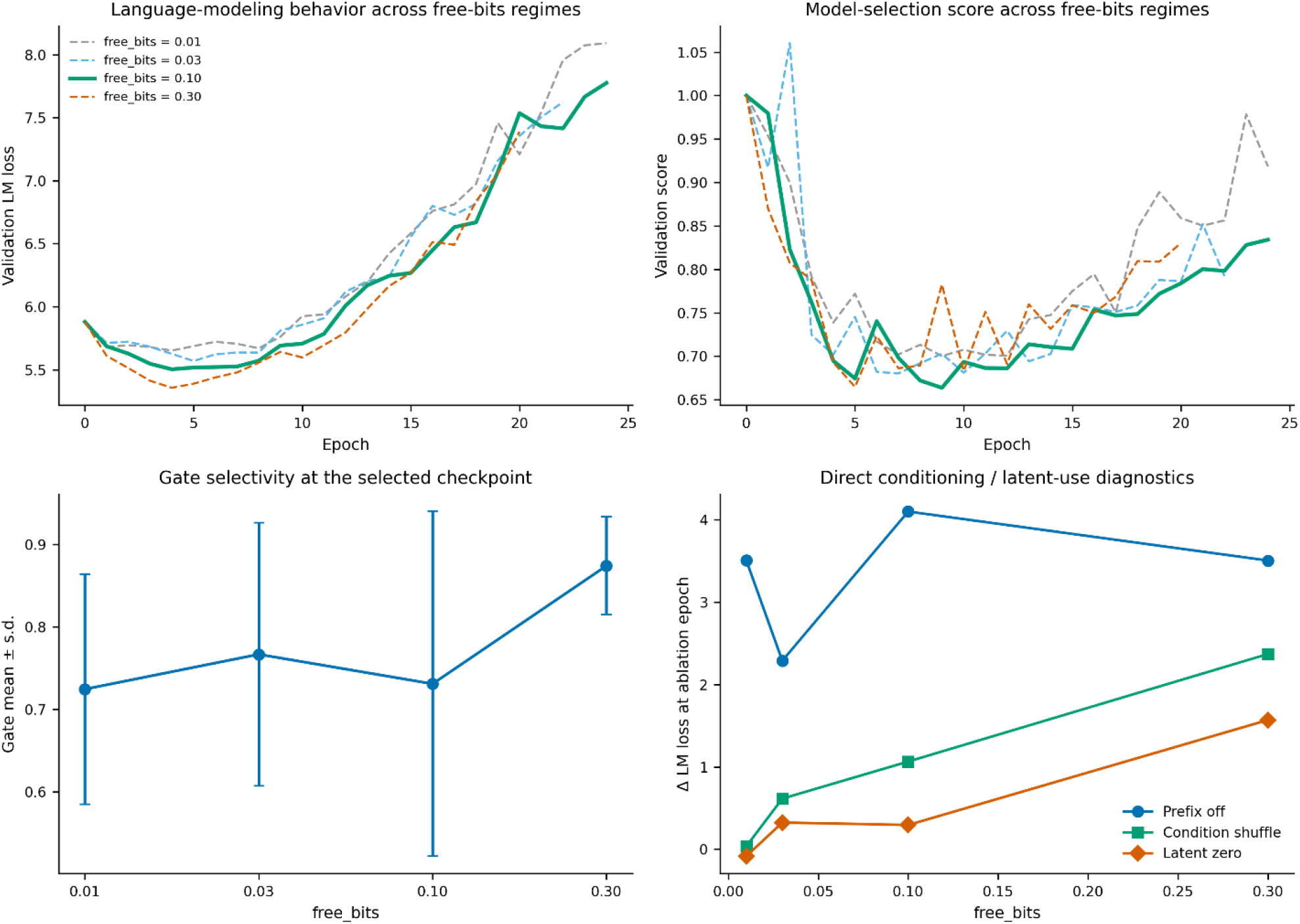
Different free bits regime evaluations. Comparison of language modeling loss, validation score, gate selectivity, and ablation experiments across four free-bits regimes, revealing a gradient in latent information flow. The optimal trade-off between regularization and generative quality is achieved at free-bits = 0.10, with higher values compressing gate dynamics and lower values rendering the prefix channel nearly inert.

When we look at the gate dynamics experiments at 0.1 free bits schedule the gate opens near 0.9, representing an early strong use of the latent. During the first 5 epochs that the gate mean remains stable, followed by a progressive reduction, achieving a mean of 0.731 at the selected epoch. As the gate means start decreasing, the standard deviation starts growing, separating the maximum from the minimum, representing an indicative of active channel utilization, spanning a range from 0.408 to 0.962.

The language and composite losses follow a similar pattern than observed previously through the rest of the regimes, confirming an expected difference between language modelling and the classification and regression heads. It is interesting to note that as the ablation diagnostics come closer to the selected epoch 9 the contribution of condition and sequence, as well as the prefix off ablation experiments values increase, while the gate mean reduces. The latent zero ablation, together with the condition shuffle remain quite stable up to the same epoch 9.

**Figure 3.**
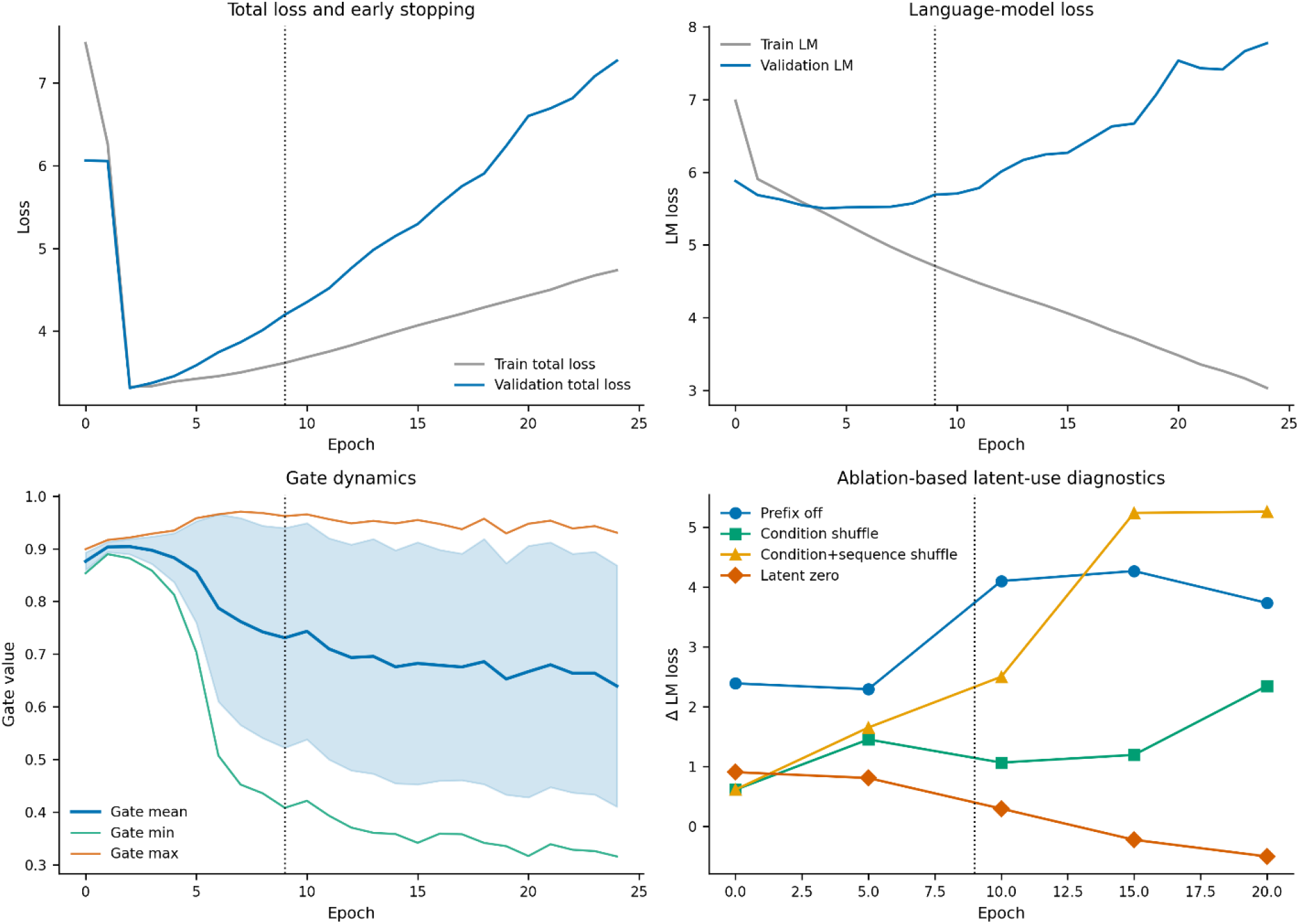
Free bits 0.1 validation analysis, gate dynamics and ablation experiments. Training and validation losses for the selected free bits = 0.10 exhibit an early optimum followed by a language modelling driven degradation. Gate dynamics reveal the transition from a uniform conditioning regime to an input-dependent variable scheme, while ablations confirm the latent relevance. The vertical dashed line marks the selected checkpoint.

### Baseline Generation: Regime Benchmarking and Bacterial Conditioning

Generation from the base CVAE was evaluated across a grid of sampling modes, perturbation magnitudes, and decoding temperatures. Two sampling modes were developed: a prior mode, in which latent vectors were drawn directly from the standard Gaussian prior, and a perturb mode, in which the encoder mean μ of a seed training sequence was used as the starting point and displaced by Gaussian noise of magnitude σ before decoding, the experimentally validated L33 sequence (KKIRVRLSA) was used all through this project to ensure reproducibility and consistency.

A Pareto Front was developed to evaluate how generation hyperparameters affect the generation and subsequently optimize the process. To construct the grid search, three different reasonable σ values were selected for each mode; 0.05, 0.20, and 0.40 for perturb mode, and 0.30, 0.50, and 0.70 for prior mode, evaluating decoding temperatures of 0.75, 0.85, and 0.95. Generated sequences were evaluated post hoc through the external ensemble scorer to classify the hit rate, diversity, and outlier propensity.

The regime selection was treated as a multi-objective optimization problem presenting three competing objectives: maximizing predicted antimicrobial quality (hit rate at MIC ≤ 1 μg/mL), maximizing sequence diversity (mean 3-mer Jaccard dissimilarity relative to the training set), and minimizing out-of-distribution risk (OOD rate). Prior and perturb modes were evaluated separately. Among the 18 unique regimes tested within both modes, 11 of them were identified as Pareto-optimal, corresponding to 61% of the explored landscape. 7 out of the 9 perturb configurations, and 4 out of 9 for prior, emerged as non-dominated regimes. One regime was selected for both generation methods (perturb regime: σ = 0.20, T = 0.95, prior regime: σ = 0.30, T = 0.95) and subsequently evaluated downstream, each prioritizing a different trade-off between potency, novelty, and physicochemical plausibility.

All perturb regimes consistently outperformed prior regimes in antimicrobial quality. The best perturb configuration (σ = 0.05, temperature = 0.75) achieved a hit rate of 42%, a mean predicted MIC of 1.09 μg/mL, a mean novelty of 0.595, and an OOD rate of 19%, while the balanced and more exploratory regime, which was selected for downstream analysis, exhibited achieved a hit rate of 38.8%, a mean predicted MIC of 1.13 μg/mL, a mean novelty of 0.668, and an OOD rate of 14%. Prior regimes produced substantially worse hit rates, ranging from 12% to 18%, exhibiting mean predicted MICs consistently above 2.0 μg/mL. The only metric in which prior generation modes surpassed perturb was the novelty, in which prior regimes yielded scores ranging from 0.691 to 0.775. It should be noted that this predicted MIC was evaluated utilizing the regression head of the decoder, which provides different values to the external score that will be discussed later in this work.

**Figure 4.**
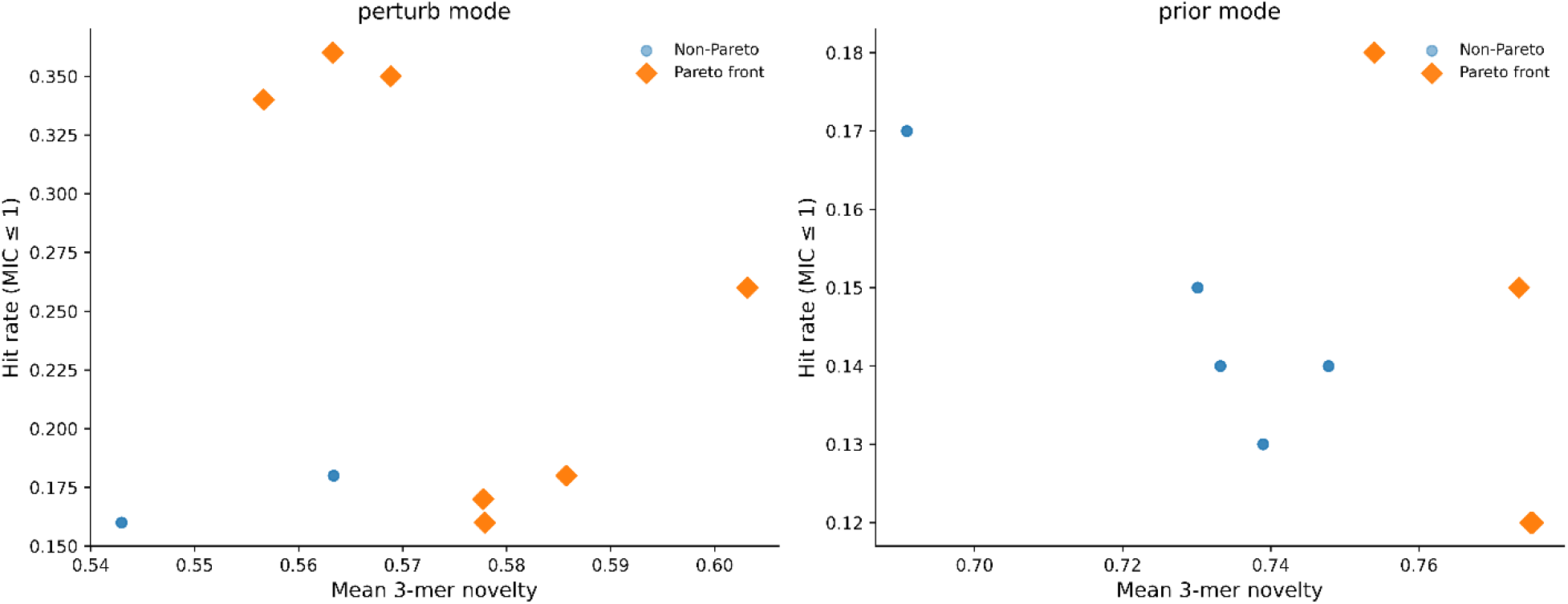
Pareto fronts by generation mode. Pareto fronts for perturb (left) and prior (right) sampling modes across hit rate at MIC ≤ 1 μg/mL and mean 3-mer novelty, with OOD rate as a third dominance criterion. Orange diamonds indicate non-dominated configurations. Perturb mode exhibits 7 out of 9 Pareto-optimal configurations while prior only exhibit 4.

### Quantitative-Structure Activity Relationship (QSAR) Guided Surrogate-Weighted Fine-Tuning (SWF) first cycle

To evaluate the generative capabilities of our CVAE we built an external ensemble composed by 5 individual Multilayer Perceptron and trained on quantitative MIC data. This new model was trained with 25 selected physicochemical characteristics selected from an initial pool of 50, it achieved a validation RMSE of 0.531 and a Spearman correlation of 0.843. The objective of this external model was to break the circularity of the CVAE and provide an external signal to guide generation implementing a more chemical point of view, which was performed through surrogate-weighted fine-tuning. This Surrogate-Weighted Fine-tuning (SWF1) was applied to the fine-tuned ProtGPT2 decoder using a pool of high-scoring candidates selected and weighted by the external QSAR ensemble. To fairly evaluate the effect of this process and determine where the effect comes from, we conducted a 3×3 factorial experiment starting from both generation regimes, including two different prior regimes. The perturb regime (σ = 0.20, T = 0.95) and the prior regime (σ = 0.30, T = 0.95) were selected from the Pareto-optimal configurations, however, the prior regime (σ = 0.70, T = 0.95) was included to evaluate how a different sigma value could condition the generation, leading into a better exploration towards boundary-proximal vectors. 500 sequences against each targeting bacteria were generated utilizing two different seeds and rescoring all outputs with a fixed external score.

The results confirmed unequivocally that the quality of the training pool used to train the ensemble is the primary determinant of output quality, even if the generation mode clearly plays a role. In perturb final regime generation, when the ensemble is trained with perturb-derived candidates (mean 1.532) the mean score outperforms the prior-trained ensemble (mean 1.762). Interestingly, both modes improve the perturb baseline, illustrating that even prior-trained ensembles provide useful signal.

The same pattern is observed during prior final regime generation, when the perturb-trained ensemble produces systematically better scores through both prior regimes (σ = 0.3, σ = 0.7). However, the prior generation at σ = 0.7 (mean score 2.03) produces significantly better candidates regime σ = 0.3 (mean score 3.16), illustrating that more explorative generation regimes benefit better from high-activity related seeds than conservative configurations. Interestingly, these results indicate that generating in prior, the SWF1 performed by prior-trained ensembles does not improve or marginally improves the quality of prior generative regimes, while SWF1 performed by perturb-trained ensembles does it.

A supplementary experiment evaluated whether Cross-entropy generated candidates could serve as a viable SWF training pool. CEM generated candidates from the baseline decoder yielded a best candidate 1.219, while exhibiting a mean score of 1.624, substantially better than the baseline decoder prior. When the ensemble is trained with this pool to reweight the decoder, and subsequently generate in perturb mode, it results into a mean score of 1.728, while generating through prior mode produces a score above 3.4. These findings confirm that the CEM strategy, although it seems to be solid for candidates generation, widely surpassing the prior decoder generation regime, fails to maintain the correct geometry needed for efficient latent reorganization.

**Figure 5.**
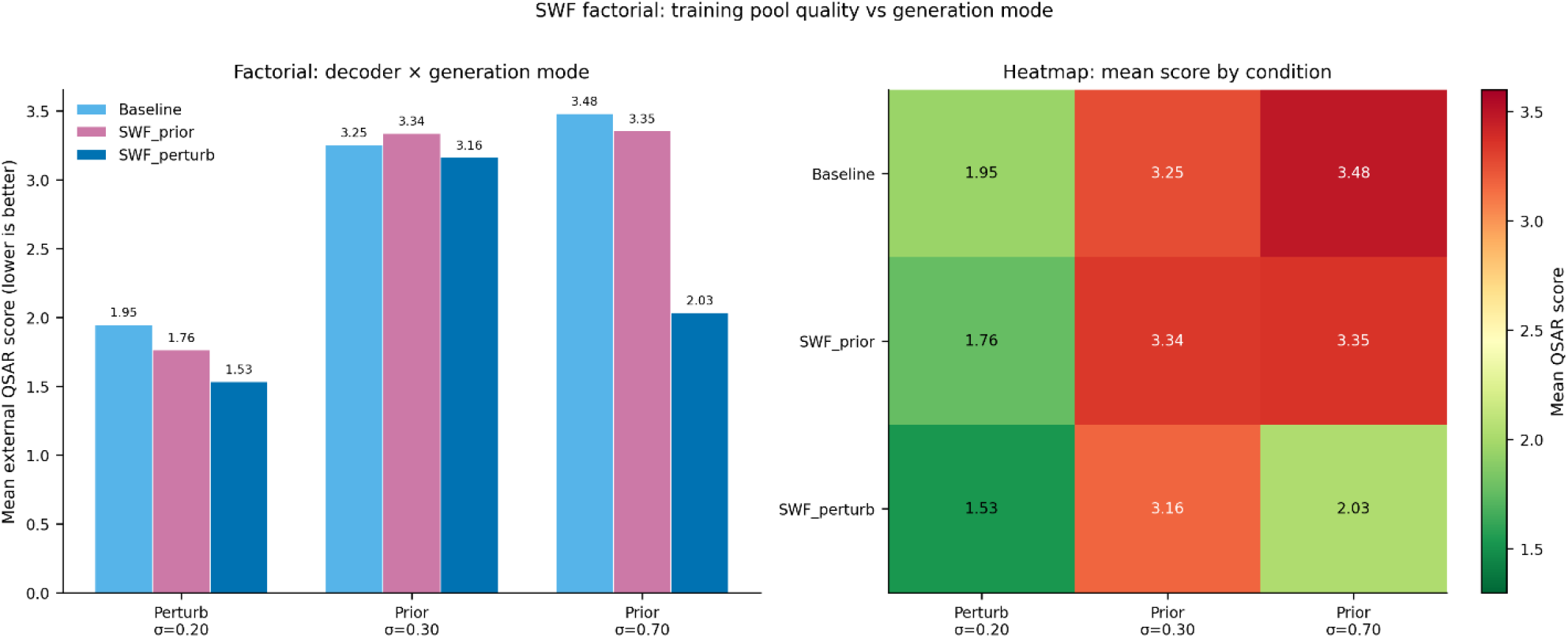
Factorial analysis of SWF training pool quality and generation mode. Left: mean external score depending on the ensemble training (baseline, SWF trained on prior, and SWF trained on perturb) and generation modes (Perturb σ=0.20, Prior σ=0.30, Prior σ=0.70). Right: heatmap corresponding to the factorial experiment.

### Latent Geometry: How SWF1 Reorganizes the Generative Distribution

CEM operates by iteratively sampling latent vectors from a Gaussian distribution, decoding them, scoring the outputs through the external ensemble, and subsequently updating the distribution towards higher-scoring regions. The success of this strategy relies on the geometry of the latent space, the smoother gradient the best exploitation. Three structural controls were evaluated by utilizing the exact same hyperparameters: the baseline, and both reweighted decoders, trained by prior and perturb regimes.

CEM applied to the baseline decoder produced a mean external score of 1.608 and a best candidate of 1.230. This represents an improvement over both the prior and perturb baselines which present scores of 1.95 and 3.25 respectively. This illustrates that CEM performs very well, while interestingly reduced the mean sequence length from 18.5 to 12.5 residues, and novelty from 0.695 to 0.521.

When the CEM strategy is applied over the reweighted decoder utilizing prior generated candidates, it produces a mean score of 2.390, presenting a best candidate of 1.448, which worsens the perturb final generation utilizing the same decoder (mean score of 1.762), but improves the prior final generation of 3.34. However, these results are at the same time substantially worse than its generation without SWF, it proves that the external physicochemical signal does not provide a better latent for sampling. Besides, this regime shows a high standard deviation (1.165) indicating that CEM finds only isolated favorable regions rather than a coherent high-activity landscape.

On the other hand, a different paradigm emerges when CEM is applied as generative sampling mode over the perturb-trained decoder, which produces a mean score of 1.398, and a best candidate of 1.191. This represents the best results across all evaluated configurations and confirms that CEM effectively samples when it starts from a reorganized latent, as the seed cluster seems to be after reweighting.

**Figure 6.**
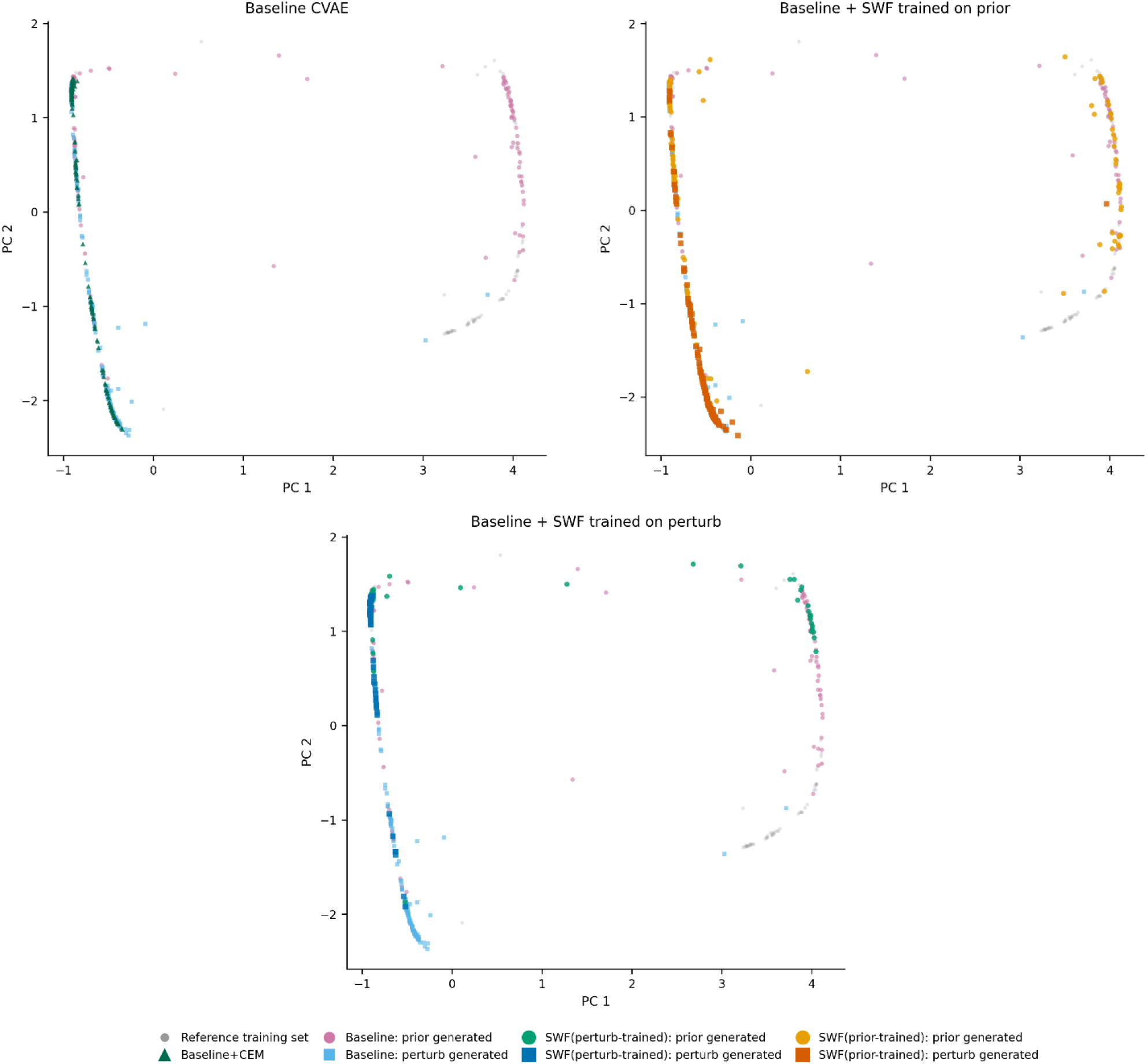
PCA of the latent space under three decoder conditions. Each panel projects 200-dimensional mean vectors onto the first two principal components fitted on the reference training set. Top left: Baseline CVAE. Top right: Baseline with SWF trained on prior sequences. Bottom: Baseline with SWF trained on perturb sequences. Surrogate weighted fine-tuning through prior generation is unable to sample close to high-activity antimicrobial peptides.

### Comparative Branches: Limits of Iterative Optimization

Cross Entropy Method has emerged as the optimal strategy for antimicrobial peptide generation, and it shines when it operates in an already reorganized latent space resulted from perturb-trained SWF. To evaluate the limits of this optimization pipeline, we performed further optimization techniques, such as additional iterative SWF cycles or reinforcement learning pipelines, attempting to further improve the best model. Both regimes have utilized the best elite buffer from the last iterative CEM cycle, this batch contained 278 unique sequences concentrated in a narrow region of the manifold.

The training of physicochemical-based ensemble corresponding to the second SWF cycle converged at epoch 2, and the lead into generation scores of 3.519, representing the worst score all through this project and occupied almost identical latent space than the reference training set. The mean cationic ratio collapsed to 0.165 and GRAVY rose to +0.138, indicating loss of the physicochemical AMP profile. Applying CEM to the top of this catastrophic forgetting partially rescued quality to a mean of 1.685, but this remained substantially worse than SWF1+CEM.

The Reinforcement Learning scheme followed was REINFORCE and established the external score as the reward function. It produced a mean score of 1.442, which was worse than the optimal configuration but avoided catastrophic degrading. Novelty increased from 0.605 to 0.662 and repetition score from 1.516 to 1.804, consistent with REINFORCE increasing sequence entropy at the cost of scoring precision.

The last experiment evaluated the generation from the decoder baseline starting from the sequences produced through the best configuration (SWF1+CEM), but the results produced a mean score of 3.242, a counterintuitive degradation explained by the fundamental mismatch between the baseline latent geometry and the high-quality anchor regions: forcing generation toward areas the model cannot decode coherently produces worse distributions than using the model’s native territory.

| Branch | Mean score | Best | Interpretation |
| --- | --- | --- | --- |
| SWF1+CEM | 1.345 | 1.161 | Recommended pipeline |
| SWF1+CEM+RL | 1.442 | 1.403 | Entropy increase, precision loss |
| SWF2 no-CEM | 3.519 | 3.327 | Catastrophic collapse |
| SWF2+CEM | 1.685 | 1.186 | Partial rescue only |
| baseline+RL (baseline anchors)+CEM | 2.993 | 1.481 | RL cannot substitute SWF |
| baseline+RL (SWF1 anchors)+CEM | 3.242 | 2.963 | Anchor mismatch degrades further |

**Figure 7.**
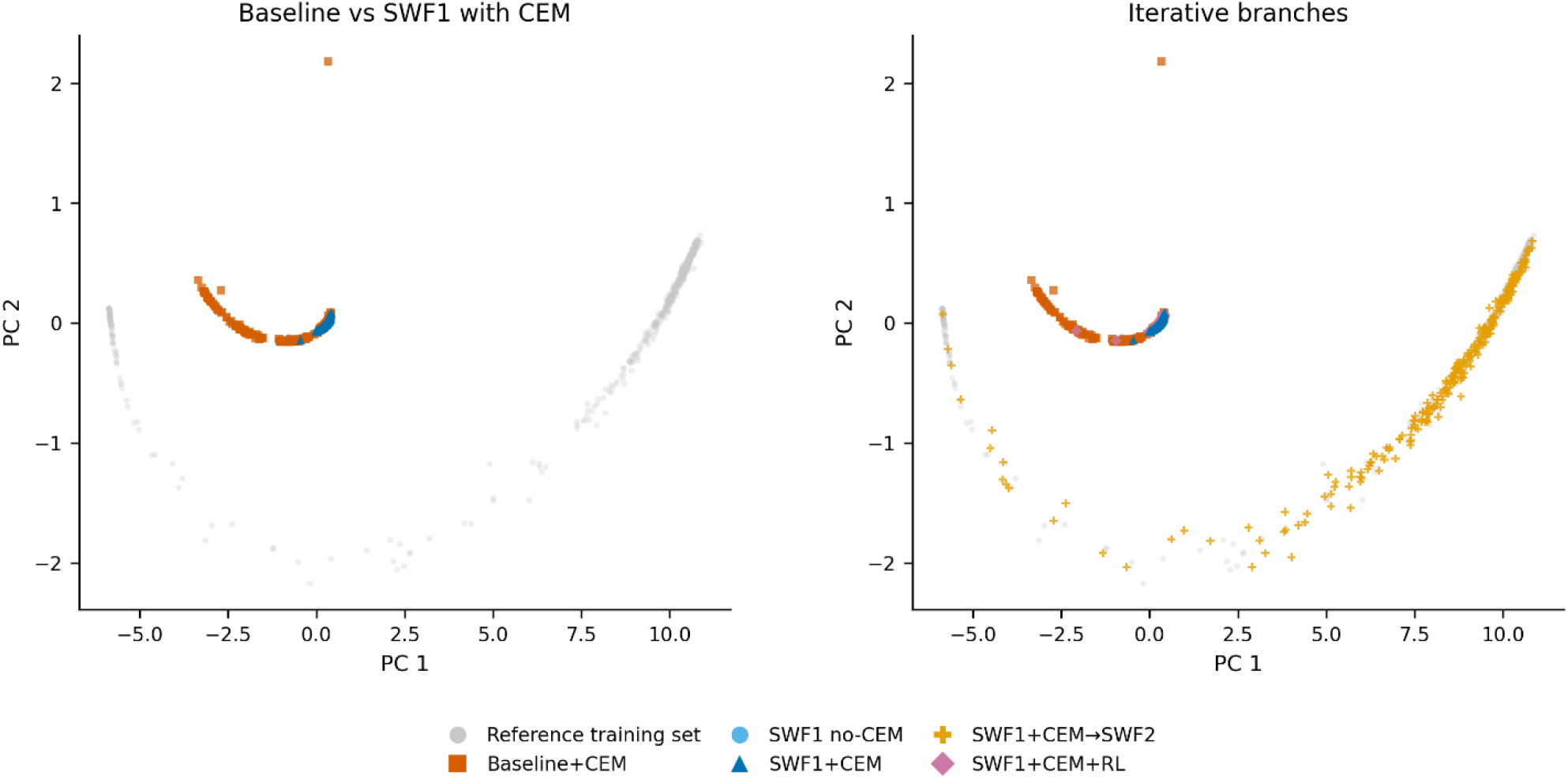
Latent space geometry of the optimal pipeline and iterative extensions. Left: PCA comparison between Cross Entropy Method generation baseline, Surrogate Weighted Fine-tuning perturb baseline, and both strategies working together, Right: Further iterative Surrogate Weighted Fine-tuning cycles and Reinforcement Learning approaches over the best-performing model.

### AMPs Generated Evaluation

To evaluate the generation of our top model we generated 500 sequences against *E. coli*, of which 465 were unique. The top 100 antimicrobial sequences by score were selected and further evaluated. The mean score was 1.259 μg/mL, with the best individual candidate scoring 1.161 μg/mL and the top ten candidates averaging 1.195 μg/mL. The mean sequence length was 17.2, and the mean net charge was 6.83, which represents a mean cationic ratio of 0.416. The mean GRAVY index was −0.421, and the hydrophobic moment was 0.598.

ESMFold was used to predict the three-dimensional structure of our top candidates, exhibiting a mean pLDDT of 83.7, above the threshold of 70 generally used to indicate a well-defined predicted structure. Analysis of backbone dihedral angles revealed that 96% of candidates exhibited helical content exceeding 0.5, with a mean helical fraction of 0.874 across the set.

To put these results in context we generated a dataset extracted from DBAASP containing 1,336 Gram-negative antimicrobial peptides presenting a length from 8 to 30 residues, and we evaluated them against our candidates. The net charge and the cationic rate of our candidates was higher than the selected peptides, as well as the hydrophobic moment. However, although our peptides candidates presented a lower standard deviation, the mean length, the GRAVY index, and the Chou-Fasman helicity score presented quite comparable values.

**Figure 8.**
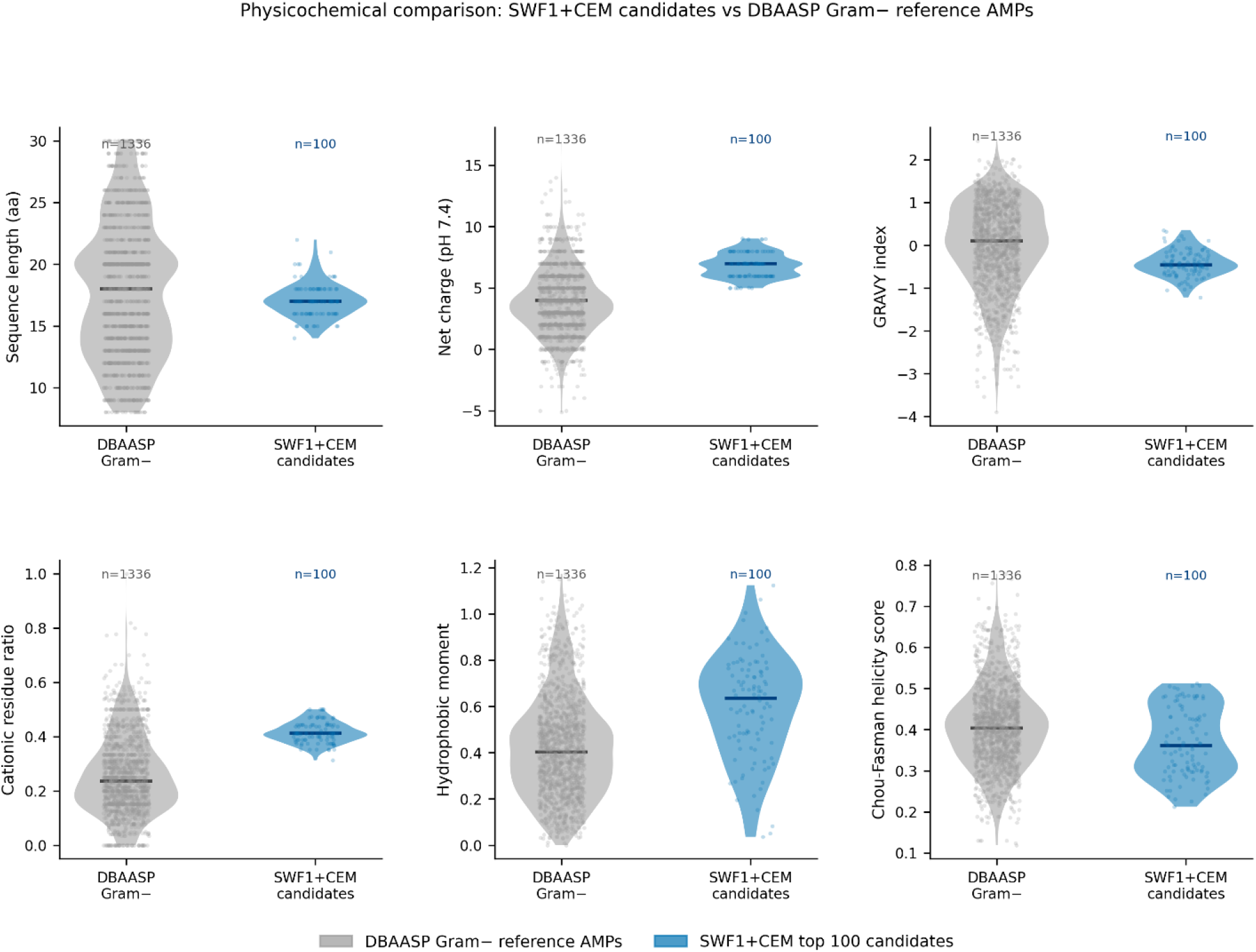
Physicochemical characterization of SWF1+CEM (best—performing model) top 100 candidates relative to DBAASP Gram-negative reference AMPs. Violin plots compare six sequence-level descriptors between top candidates and 1,336 experimentally validated Gram-negative AMPs from the DBAASP database. Horizontal lines indicate medians. Individual data points are shown as jittered scatter.

To provide an independent computational validation, our model was also evaluated against APEX model from the Machine Biology group of the University of Penn^17^, a multitask deep learning model trained on experimental MIC data using a distinct architecture and training dataset. Four different regimes were evaluated: the three baselines (prior, perturb and CEM), as well as the best model configuration. The best regime (SWF1+CEM) exhibited lower mean predicted MIC than all three baseline regimes across the three target organisms: 34.4 vs 44.4, 37.1, and 54.6 μg/mL for *E. coli*; 132.9 vs 184.6, 186.2, and 176.8 μg/mL for *K. pneumoniae*; and 59.5 vs 74.9, 71.0, and 87.1 μg/mL for *P. aeruginosa* (quality-first, balanced, and exploration respectively), proving the efficacy of our ensemble independently. The Spearman correlation between the internal QSAR scores and APEX predictions was low across all three organisms (r = 0.06, 0.21, and −0.09 for *E. coli, K. pneumoniae, and P. aeruginosa*), confirming that the two scorers are effectively independent.

**Figure 9.**
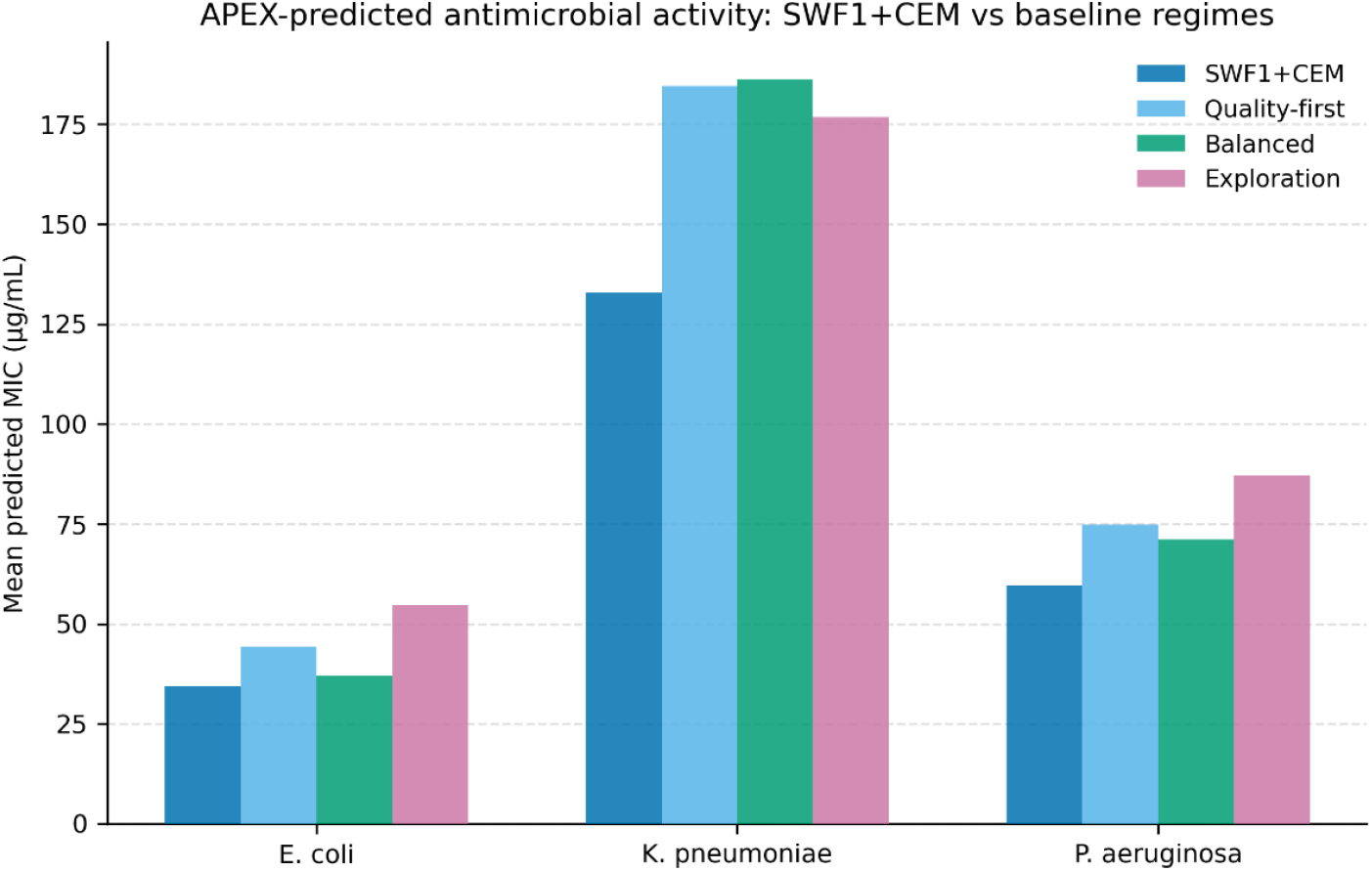
Independent APEX validation of SWF1+CEM candidates relative to baseline regimes. Mean predicted MIC from the APEX model across the three target organisms. SWF1+CEM candidates consistently exhibit lower predicted MIC than all baseline regimes, providing independent validation from a model architecturally distinct from the internal QSAR ensemble.

## Discussion

The pipeline presented in this work rests on a solid architectural foundation. The transformer-based encoder successfully developed a discriminative latent space organized depending on binary antimicrobial classification (AMP/noAMP), exhibiting AUROC values of 0.968 while utilizing 61.1 out of 64 dimensions. A gating scalar function has been implemented to control the magnitude of the latent flow towards the decoder; ProtGPT2, a LLM model trained on protein sequences.^26^ This decoder was fine-tuned through Low-Rank Adaptation (LoRA) technique utilizing a quantitative bacteria specific dataset, expressing efficacy as MIC value in logarithmic scale. Gate dynamics and ablation experiments confirmed that the gating mechanism was able to regularize the latent channel adjusting to decoder requirements dynamically through a scalar function, exhibiting prefix removal degrading language modeling loss by 4.10 nats and bacterial condition shuffling producing a degradation of 1.07 nats, which seemed to be the most effective ablations experiments to evaluate the distinct free bits regimes. These findings illustrate that both the prefix mechanism and the bacterial label contribute meaningfully to generation.

The species-specific fine-tuning strategy proved to be ineffective to achieve its objective, generated sequences against all the bacteria demonstrated to show matched-regime Jaccard similarity of 1.0. The bacteria label token produced non-zero logit perturbations at the decoder input level, but apparently not sufficiently potent to learn a discriminative effect. All downstream experiments were subsequently performed for *E. coli*.

The generator was set up in two different modes; prior, perturb. Both were evaluated through Pareto front construction, proposed as a multiobjective problem, aiming to maximize predicted antimicrobial quality (hit rate at MIC ≤ 1 μg/mL), maximize sequence diversity (mean 3-mer Jaccard dissimilarity relative to the training set), and minimize out-of-distribution risk (OOD rate). The best configurations were then compared. The perturb mode, which used the experimentally validated linear sequence L33 (KKIRVRLSA) as a seed, demonstrated to outperform the best prior configuration constantly, which proved to be more effective for exploration. This result is unequivocally closely linked to the initial seed utilized.

Given that the CVAE proved to be stable and efficient to generate reasonable well-shaped antimicrobial sequences, we developed a second phase focused on constructing an ensemble MLP module based on physicochemical characteristics. This new model was trained with the same quantitative dataset utilized for the decoder module, implementing 25 physicochemical descriptors selected by mRMR from an initial pool of 50. The objective of this module was to act as an external scorer, independent from the core generator, providing conservative upper-confidence-bound predictions that penalize uncertain candidates. This fine-tuning was performed through reweighting of the decoder utilizing the best-scoring candidates from each generation mode.

To evaluate the role of the external score, we designed a factorial experiment evaluating both prior- and perturb-trained ensembles, as well as the baseline decoder, through three different generations; prior (σ = 0.3, σ = 0.7) and perturb (σ = 0.3). This produced interesting results following the line of the already commented results; 1) The perturb-trained ensemble surpassed the prior-trained ensemble at every experiment. 2) When the prior-trained ensemble is used to fine tune the decoder, and the final generation mode is also prior, the results are quite poor. 3) A prior-trained ensemble led the decoder to generate better candidates in perturb generation mode than the perturb baseline, which means that the prior candidates contain valuable signal codified as physicochemical properties. 4) A perturb-trained ensemble can lead to poor, or reasonably good generation depending on the sigma value set up when synthesizing in prior mode, and a sigma of 0.7 generates scores worse, but close to the perturb baseline. All these findings support the idea that enrich the general prior and generating close to high-active clusters is a viable strategy despite the prior generation remains weak.

In a secondary phase, we utilized Cross Entropy Methodology to test if this could beat the minimal prior generation through evolutionary sampling, based on the external score. CEM proved to generate better than both previous generation regimes; prior and perturb, emerging as the top baseline generation mode (mean score 1.608). CEM was also evaluated over two different decoders; prior-trained ensemble fine-tuned and perturb-trained ensemble fine-tuned, and improved both regimes, achieving the best model score for perturb-trained ensemble fine-tuned exhibiting a score of 1.398. However, it is interesting to note that CEM baseline generated candidates were not useful for SWF reweighting, this likely means that this method performs as a great sampler, but the candidates come from so different latent spaces that are not able to produce an smooth gradient for subsequent generations.

Further iterative ensemble-based fine tuning was explored, as well as Reinforcement Learning approaches, but none of them surpassed the best model. A second iterative cycle resulted in catastrophic degradation of the decoder weights, as can be observed in the UMAP analysis, while RL maintained most of its efficacy, but was unable to improve it, neither at the top of the best model, or working over the decoder baseline.

A pool of one hundred peptides was generated by the best model achieved, and their mean physicochemical properties were analyzed, and subsequently compared against a dataset of Gram-negative antimicrobial peptides. Similar characteristics were found, despite the net charge, and the hydrophobic moment, which seemed to be higher in our synthetic peptides. Besides, all the characteristics presented lower standard deviation in our studies. A structural analysis of our peptides was also performed utilizing ESMFold,^33^ indicating that 96% of candidates exhibited helical content exceeding 0.5, with a mean helical fraction of 0.874 across the set, besides, the pool exhibited a mean pLDDT of 83.7, above the threshold of 70 generally used to indicate a well-defined predicted structure. These findings indicate that our model was able to generate chemically viable, novel, and stable molecules.

APEX model, property of the Machine Learning Group of the University of Pennsylvania was used for external validation,^17^ and four of our models were evaluated; prior, perturb, and CEM baseline, as well as the best scoring model. The best scoring model achieved to obtain better efficacy against all the targeted bacteria: *E. coli, P. aeruginosa* and *K. pneumoniae*, externally proving the success of our methodology.

## Conclusion

This work presents a modular generative pipeline for targeted antimicrobial peptide design that addresses two fundamental limitations: the scarcity and quality of experimentally validated data, and the circular dependency between generation and evaluation. The transformer-based encoder succeeds in generating a discriminative 64-dimensional latent space, exhibiting AUROC 0.968, F1 0.919, while retaining 61.1/64 active dimensions. A fine-tuned ProtGPT2 was utilized as decoder, leveraging latent information through gated soft-prompting, and a free bits optimization strategy, which yielded well-regularized latent and competitive early candidates. The use of an external ensemble QSAR-based proved to be a successful strategy to break the circularity and guide the exploration and exploitation within high activity clusters, which was supported by external validation utilizing APEX model, one of the field’s leading independent scoring models. The Cross-Entropy Method demonstrated the ability to generate promising candidates through iterative latent sampling without requiring experimentally validated high-activity seeds, and both strategies proved to efficiently work together, yielding the best-performing model, which was ultimately utilized to generate the definitive pool of peptide candidates. These peptides exhibited physicochemical characteristics consistent with Gram-negative antimicrobial peptides reported in the literature, alongside significant helical content (mean helical fraction of 0.874) and high-confidence structural predictions (mean pLDDT 83.7).

Some limitations were identified during this work; although the decoder was trained in a species-specific way, this did not translate into discriminative generation across bacterial targets. Furthermore, experimental validation will be necessary to establish the biological relevance of the generated candidates.

## Contributions

I.C. conceived the study, designed the methodology, curated the datasets, developed the computational pipeline, trained and evaluated the models, performed the analyses, prepared the figures, and wrote the original draft of the manuscript. F.W., C.F.N., C.F., and A.P. contributed to the critical revision of the manuscript and provided feedback on the interpretation and presentation of the results. All authors reviewed and approved the final version of the manuscript. C.F., and A.P. contributed equally to this work.

## Code availability

The code used in this study is publicly available at Zenodo: 10.5281/zenodo.19697476. The corresponding source repository is available on GitHub: Ismaelcasku/cvae-protgpt2-amp.

## Funding Sources

This work was supported by a European Union Horizon Europe MSCA DN-ID grant (grant number 101073263). https://ssbb-project.eu/.

## Notes

https://zenodo.org/records/19697476

### Competing Interest Statement

The authors have declared no competing interest.

### Summary of Updates

The name of author number 3 was corrected.

## References

1. Magana, M. et al. The value of antimicrobial peptides in the age of resistance. The Lancet Infectious Diseases 20, e216–e230 (2020). 10.1016/S1473-3099(20)30327-3

2. Singh, A., Tanwar, M., Singh, T. P., Sharma, S. & Sharma, P. An escape from ESKAPE pathogens: A comprehensive review on current and emerging therapeutics against antibiotic resistance. International journal of biological macromolecules 279, 135253 (2024). 10.1016/j.ijbiomac.2024.135253

3. Huan, Y., Kong, Q., Mou, H. & Yi, H. Antimicrobial Peptides: Classification, Design, Application and Research Progress in Multiple Fields. Frontiers in Microbiology 11, 582779 (2020). 10.3389/fmicb.2020.582779

4. Talapko, J. et al. Antimicrobial Peptides—Mechanisms of Action, Antimicrobial Effects and Clinical Applications. Antibiotics 11, 1417 (2022). 10.3390/antibiotics11101417

5. Yin, L. M., Edwards, M. A., Li, J., Yip, C. M. & Deber, C. M. Roles of hydrophobicity and charge distribution of cationic antimicrobial peptides in peptide-membrane interactions. Journal of Biological Chemistry 287, 7738–7745 (2012). 10.1074/jbc.M111.303602

6. Zheng, S. et al. Antimicrobial peptide biological activity, delivery systems and clinical translation status and challenges. Journal of Translational Medicine 23, 292 (2025). 10.1186/s12967-025-06321-9

7. Gagat, P., Ostrówka, M., Duda-Madej, A. & Mackiewicz, P. Enhancing Antimicrobial Peptide Activity through Modifications of Charge, Hydrophobicity, and Structure. International Journal of Molecular Sciences 25, 10821 (2024). 10.3390/ijms251910821

8. AL Khathami, M. M. M., Alenazi, A. M., Alareefi, H. S. & Alomran, R. W. Understanding antibiotic resistance: Challenges and solutions. Int. J. Health Sci. (Qassim). 5, 1255–1274 (2021).

9. Nordström, R. & Malmsten, M. Delivery systems for antimicrobial peptides. Adv. Colloid Interface Sci. 242, 17–34 (2017). 10.1016/j.cis.2017.01.005

10. Imura, Y., Nishida, M. & Matsuzaki, K. Action mechanism of PEGylated magainin 2 analogue peptide. Biochim. Biophys. Acta Biomembr. 1768, 2578–2585 (2007). 10.1016/j.bbamem.2007.06.013

11. Merz, M. L. et al. De novo development of small cyclic peptides that are orally bioavailable. Nat. Chem. Biol. 20, 624–633 (2024). 10.1038/s41589-023-01496-y

12. Martian, P. C. et al. Cyclic peptides: A powerful instrument for advancing biomedical nanotechnologies and drug development. Journal of Pharmaceutical and Biomedical Analysis 252, 116488 (2025). 10.1016/j.jpba.2024.116488

13. Bellotto, O. et al. Polymer Conjugates of Antimicrobial Peptides (AMPs) with D-Amino Acids (D-aa): State of the Art and Future Opportunities. Pharmaceutics 14, 446 (2022). 10.3390/pharmaceutics14020446

14. Dean, S. N. & Walper, S. A. Variational autoencoder for generation of antimicrobial peptides. ACS Omega 5, 20746–20754 (2020). 10.1021/acsomega.0c00442

15. Van Oort, C. M., Ferrell, J. B., Remington, J. M., Wshah, S. & Li, J. AMPGAN v2: Machine Learning-Guided Design of Antimicrobial Peptides. J. Chem. Inf. Model. 61, 2198–2207 (2021). 10.1021/acs.jcim.0c01441

16. Wang, J. et al. Discovery of Antimicrobial Peptides with Notable Antibacterial Potency by an LLM-Based Foundation Model. Sci. Adv. 11, eads8932 (2025). https://www.science.org/doi/10.1126/sciadv.ads8932

17. Torres, M. D. T. et al. A generative artificial intelligence approach for antibiotic optimization. Preprint at 10.1101/2024.11.27.625757 (2024).

18. Hu, E. J. et al. LoRA: Low-Rank Adaptation of Large Language Models. Int. Conf. Learn. Represent. (2022).

19. Srinivas, N., Krause, A., Kakade, S. M. & Seeger, M. W. Information-Theoretic Regret Bounds for Gaussian Process Optimization in the Bandit Setting. IEEE Trans. Inf. Theory 58, 3250–3265 (2012).

20. Lakshminarayanan, B., Pritzel, A. & Blundell, C. Simple and Scalable Predictive Uncertainty Estimation using Deep Ensembles. Adv. Neural Inf. Process. Syst. 30 (2017).

21. De Boer, P.-T., Kroese, D. P. & Rubinstein, R. Y. A Tutorial on the Cross-Entropy Method. Ann. Oper. Res. 134, 19–67 (2005).

22. Devlin, J., Chang, M.-W., Lee, K. & Toutanova, K. BERT: Pre-Training of Deep Bidirectional Transformers for Language Understanding. Proc. 2019 Conf. North American Chapter Assoc. Comput. Linguistics 4171–4186 (2019).

23. Vaswani, A. et al. Attention Is All You Need. Adv. Neural Inf. Process. Syst. 30 (2017).

24. He, J., Spokoyny, D., Neubig, G. & Berg-Kirkpatrick, T. Lagging Inference Networks and Posterior Collapse in Variational Autoencoders. Int. Conf. Learn. Represent. (2019).

25. Steinegger, M. & Söding, J. MMseqs2 enables sensitive protein sequence searching for the analysis of massive data sets. Nat. Biotechnol. 35, 1026–1028 (2017).

26. Ferruz, N., Schmidt, S. & Höcker, B. ProtGPT2 is a deep unsupervised language model for protein design. Nat. Commun. 13, 4348 (2022). 10.1038/s41467-022-32007-7

27. Li, X. L. & Liang, P. Prefix-Tuning: Optimizing Continuous Prompts for Generation. Proc. 59th Annu. Meet. Assoc. Comput. Linguistics 4582–4597 (2021).

28. Deb, K., Pratap, A., Agarwal, S. & Meyarivan, T. A Fast and Elitist Multiobjective Genetic Algorithm: NSGA-II. IEEE Trans. Evol. Comput. 6, 182–197 (2002).

29. Rotem, S., Radzishevsky, I. & Mor, A. Physicochemical properties that enhance discriminative antibacterial activity of short dermaseptin derivatives. Antimicrob. Agents Chemother. 50, 2666–2672 (2006).

30. Özçelik, R., van Weesep, L., de Ruiter, S. & Grisoni, F. peptidy: a light-weight Python library for peptide representation in machine learning. Bioinformatics Advances 5, vbaf058 (2025). 10.1093/bioadv/vbaf058

31. Peng, H., Long, F. & Ding, C. Feature Selection Based on Mutual Information: Criteria of Max-Dependency, Max-Relevance, and Min-Redundancy. IEEE Trans. Pattern Anal. Mach. Intell. 27, 1226–1238 (2005).

32. Zhang, J., Kim, J., O’Donoghue, B. & Boyd, S. Sample Efficient Reinforcement Learning with REINFORCE. Int. Conf. Learn. Represent. (2021).

33. Lin, Z. et al. Evolutionary-scale prediction of atomic-level protein structure with a language model. Science 379, 1123–1130 (2023). https://www.science.org/doi/10.1126/science.ade2574

