## Supplementary Materials for "A Conditional Variational Autoencoder with QSAR-Guided Surrogate-Weighted Fine-Tuning and Cross-Entropy Optimization for Targeted Antimicrobial Peptide Generation"

#### Supplementary Methods

##### SM1. Encoder Architecture and Training Hyperparameters

The encoder implemented a transformer-based architecture utilizing CLS token pooling strategy. The classification head operates on the sampled vector z during training, while the deterministic posterior mean μ is used for downstream conditioning and latent-space analyses. Free-bits strategy was evaluated and proved useful to prevent posterior collapse. Moreover, word-dropout was implemented to reduce over-fitting.

Table SM1a reports the architecture hyperparameters and Table SM1b reports the training configuration. The dataset comprised 5,000 sequences with binary AMP/noAMP labels partitioned at the MMseqs2 cluster level (identity 0.70, coverage 0.80) into 3,483 training, 776 validation, and 741 test sequences.

***Table SM1a. Encoder architecture hyperparameters.***

| Parameter | Value |
|---|---|
| Architecture | Transformer encoder (batch_first) |
| Number of layers | 4 |
| Attention heads | 8 |
| Model dimensionality (d_model) | 256 |
| Feed-forward dimensionality | 1024 |
| Latent dimensionality | 64 |
| Dropout | 0.10 |
| Max sequence length | 64 tokens |
| Positional encoding | Sinusoidal, masked on padding |
| Vocabulary size | 24 tokens (20 AA + PAD, MASK, CLS, EOS) |
| Pooling strategy | CLS token representation |
| Classification head input | Sampled z (training); μ used for CVAE conditioning |

***Table SM1b. Encoder training hyperparameters.***

| Parameter | Value |
| --- | --- |
| Training epochs (maximum) | 20 |
| Early stopping criterion | Validation BCE loss (minimize) |
| Selected checkpoint | Epoch 9 (val AUROC 0.949) |
| Optimizer | AdamW |
| Learning rate | $1 \times 10^{-3}$ |
| KL regularization | Free-bits (minimum rate = 0.10 nats/dim) |
| Word dropout rate | 0.10 |
| Loss function | BCE (classification) + KL divergence |
| Dataset (train / val / test) | 3,483 / 776 / 741 sequences |

#### SM2b. CVAE Base Model Architecture and Training Hyperparameters

The CVAE full architecture is the core generator of the pipeline, and it connects the encoder to ProtGPT2, an LLM trained on 700M parameters to generate proteins. ProtGPT2 decoder was fine-tuned via LoRA, and received the latent information regularized though a gated prefix conditioning mechanism. To connect the 64-dimensional latent space to the 768 dimensions of ProtGPT2 we utilized a two-layers MLP.

A scalar gate

$$g = g_{\min} + (1 - g_{\min})\ \mathrm{sigmoid}\,(W_g + b_g)$$

modulates the prefix magnitude. The full hyperparameter configuration is listed below.

***Table SM2b. CVAE base model hyperparameters.***

| Parameter | Value |
|---|---|
| Decoder base model | ProtGPT2 (nferruz/ProtGPT2, 36 layers, 738M parameters) |
| Fine-tuning strategy | LoRA (Low-Rank Adaptation) |
| Latent dimensionality | 64 |
| Prefix length | 16 tokens |
| Gate floor (g_min) | 0.2 |
| Gate input | Posterior mean μ (deterministic) |
| Encoder status during training | Frozen (freeze_encoder_epochs = 0, trained from epoch 0) |
| Free-bits (selected) | 0.10 nats/dimension |
| β schedule | Cyclical annealing, 0 → 1 over training |
| Learning rate | $2 \times 10^{-4}$ |
| Weight decay | 0.01 |
| Maximum epochs | 120 |
| Batch size | 16 |
| Selected checkpoint | Epoch 9, val composite score 0.664 |
| Precision | bf16-mixed |
| Bacterial conditions | E. coli (<ECOLI>), K. pneumoniae (<KPN>), P. aeruginosa (<PAER>) |
| Training dataset (rows / unique seqs) | 7,638 / 2,546 |
| Validation dataset (rows / unique seqs) | 849 / 283 |

#### SM2. CVAE Decoder Training Dataset

The decoder training dataset comprises 2829 non-redundant antimicrobial peptides from DBAASP, DRAMP, APD3, and CAMPR4. The activity of the peptides is expressed as MIC, corresponding to their experimentally validated values (log-transformed µg/mL). The negative dataset was built utilizing UniProt sequences, extracted from random proteins and validated as non-antimicrobial through CAMPR4 and DRAMP predictors. The negative sequences exhibited similar length distribution to average AMPs, and were partitioned following the same MMseqs2-based cluster-level strategy described in the Methods (identity threshold 0.70, coverage 0.80), yielding 2,546 training and 283 validation unique sequences

***Table SM2. CVAE decoder dataset composition by organism.***

| Organism / Set | Sequences (n) | Label type | Data sources |
|---|---|---|---|
| E. coli | 624 | MIC (log µg/mL) | DBAASP, DRAMP, APD3, CAMPR4 |
| P. aeruginosa | 602 | MIC (log µg/mL) | DBAASP, DRAMP, APD3, CAMPR4 |
| K. pneumoniae | 261 | MIC (log µg/mL) | DBAASP, DRAMP, APD3, CAMPR4 |
| Negative (all) | 1,512 | MIC = 4.0 (log) | DBAASP, DRAMP, APD3, CAMPR4 |
| Total unique | 2,829 | — | — |

#### SM3. QSAR Ensemble: Architecture, Feature Selection, and Per-Seed Validation

The external QSAR ensemble is composed of 5 individual HeteroHLP models utilizing different random seeds for each of them. The models contain three hidden layers of 256 units, implementing LayerNorm and GELU activation scheme. Since the models are trained on quantitative MIC data (log-transformed μg/mL), a heteroscedastic output head compute the uncertainty term by predicting both mean and log-variance of the predicted MIC distribution. 25 physicochemical features were utilized to train the models, selected from a initial pool of 50, computed by Peptidy library at pH 7.0. Features correlated with sequence length (Pearson |r| > 0.7) were normalized by length prior to selection.

***Table SM3a. Per-seed validation metrics for the QSAR ensemble (n = 5 seeds).***

| Seed | Best val. NLL | Val. RMSE | Val. Spearman ρ |
|---|---|---|---|
| 1 | 0.343 | 0.531 | 0.846 |
| 2 | 0.402 | 0.553 | 0.843 |
| 3 | 0.495 | 0.527 | 0.841 |
| 4 | 0.471 | 0.530 | 0.844 |
| 5 | 0.491 | 0.534 | 0.840 |
| Ensemble | — | 0.531 | 0.843 |

The 25 features retained by mRMR are listed below:

***Table SM3b. Physicochemical features selected by mRMR (sorted by selection order).***

| # | Feature name | # | Feature name |
|---|---|---|---|
| 1 | isoelectric_point | 14 | freq_T |
| 2 | charge | 15 | freq_D |
| 3 | charge_density | 16 | freq_R |
| 4 | n_H | 17 | freq_W |
| 5 | n_h_acceptors | 18 | freq_Q |
| 6 | freq_K | 19 | freq_S |
| 7 | n_N | 20 | freq_N |
| 8 | n_C | 21 | aliphatic_index |
| 9 | freq_E | 22 | energy_based_on_logP |
| 10 | average_number_rotatable_bonds | 23 | aromaticity |
| 11 | n_O | 24 | freq_P |
| 12 | molecular_weight | 25 | freq_Y |
| 13 | freq_M | — | — |

#### SM4. Free-Bits Regime Comparison

Four free-bits schemes were evaluated to infer the optimal regime, balancing meaning and regularization. Regimes under 0.1 reflected to be so regularized to be informative, but higher free bits values of 0.1-0.3 dilute the signal.

***Table SM4a. Best-epoch validation metrics across four free-bits regimes.***

| Free-bits | Best epoch | Val. score | Val. LM loss | Val. KL (nats) | Gate mean | Gate std | Gate range |
|---|---|---|---|---|---|---|---|
| 0.01 | 9 | 0.700 | 5.762 | 0.348 | 0.724 | 0.139 | 0.518–0.893 |
| 0.03 | 7 | 0.680 | 5.637 | 1.336 | 0.767 | 0.159 | 0.516–0.942 |
| 0.10 (selected) | 9 | 0.664 | 5.692 | 4.347 | 0.731 | 0.209 | 0.408–0.962 |
| 0.30 | 5 | 0.665 | 5.389 | 11.570 | 0.874 | 0.059 | 0.788–0.952 |

***Table SM4b. Ablation diagnostics at nearest available epoch (Δ in language modeling loss, nats).***

| Free-bits | Δ prefix removed | Δ condition shuffled | Δ condition + seq. shuffled | Δ latent zeroed |
|---|---|---|---|---|
| 0.01 | 3.508 | 0.036 | 1.452 | −0.084 |
| 0.03 | 2.290 | 0.615 | 0.873 | 0.324 |
| 0.10 (selected) | 4.102 | 1.065 | 2.500 | 0.295 |
| 0.30 | 3.507 | 2.371 | 2.406 | 1.570 |

*† Negative Δ indicates that the ablated condition produced lower loss than the conditioned baseline, consistent with the latent being unused (free_bits = 0.01) or the model exploiting the shuffled condition as noise regularization.*

#### SM5. Base Generation Grid: Full Regime Summary

A pareto front was developed for both generation regimes; prior and perturb, evaluating 18 regimes for each bacterial target across two independent random seeds. The pareto front was stablished as a multiobjective problem which tries to optimize temperature and sigma while minimizing hit-rate (MIC ≤ 1 µg/mL), maximizing novelty (3-mer Jaccard dissimilarity ) and maximizing OOD (fraction of sequences failing hard physicochemical constraints; length 8–25, cationic ratio 0.15–0.55, net charge 0–12, GRAVY −1.5 to 2.5).

★ marks regimes selected for downstream experiments.

***Table SM5. Generation grid results for the base CVAE (E. coli, pooled across seeds).***

| Mode | σ | T | Hit rate | Mean MIC† | Novelty | OOD rate | Pareto-optimal |
|---|---|---|---|---|---|---|---|
| Perturb | 0.05 | 0.75 | 36.0% | 1.169 | 0.563 | 17.0% | Yes |
| Perturb | 0.05 | 0.85 | 16.0% | 1.350 | 0.543 | 18.0% | No |
| Perturb | 0.05 | 0.95 | 16.0% | 1.315 | 0.578 | 14.0% | Yes |
| Perturb | 0.20 | 0.75 | 35.0% | 1.184 | 0.569 | 18.0% | Yes |
| Perturb | 0.20 | 0.85 | 18.0% | 1.319 | 0.563 | 16.0% | No |
| Perturb | 0.20 | 0.95 | 18.0% | 1.322 | 0.586 | 15.0% | Yes ★ |
| Perturb | 0.40 | 0.75 | 34.0% | 1.188 | 0.557 | 16.0% | Yes |
| Perturb | 0.40 | 0.85 | 26.0% | 1.292 | 0.603 | 21.0% | Yes |
| Perturb | 0.40 | 0.95 | 17.0% | 1.330 | 0.578 | 14.0% | Yes |
| Prior | 0.30 | 0.75 | 17.0% | 2.059 | 0.691 | 21.0% | No |
| Prior | 0.30 | 0.85 | 15.0% | 2.021 | 0.730 | 21.0% | No |
| Prior | 0.30 | 0.95 | 15.0% | 2.074 | 0.773 | 22.0% | Yes ★ |
| Prior | 0.50 | 0.75 | 14.0% | 2.216 | 0.733 | 27.0% | No |
| Prior | 0.50 | 0.85 | 13.0% | 2.148 | 0.739 | 24.0% | No |
| Prior | 0.50 | 0.95 | 12.0% | 2.225 | 0.775 | 20.0% | Yes |
| Prior | 0.70 | 0.75 | 18.0% | 2.212 | 0.754 | 21.0% | Yes |
| Prior | 0.70 | 0.85 | 14.0% | 2.302 | 0.748 | 26.0% | No |
| Prior | 0.70 | 0.95 | 12.0% | 2.506 | 0.775 | 24.0% | Yes |

*† Predicted MIC values are from the decoder regression head, not from the external QSAR ensemble. External QSAR scores are reported in the main text.*

#### SM6. Bacterial Conditioning Check

50 antimicrobial sequences against all three bacterial targets were evaluated through jaccard similarity of 3-mer to infer whether the species-specific LoRA fine-tuning was able to discriminate between species. The results reflected that while non-zero logit perturbations were found at decoder input level, these perturbations were insufficient to propagate into distinct output distributions.

***Table SM6. Bacterial conditioning verification: pairwise Jaccard similarity and metric comparison across conditions.***

| Comparison | Regime | 3-mer Jaccard sim. | Mean MIC (left / right) | Novelty (left / right) |
|---|---|---|---|---|
| E. coli vs. K. pneumoniae | Perturb σ=0.20, T=0.95 | 1.000 | 1.169 / 1.169 | 0.569 / 0.569 |
| E. coli vs. P. aeruginosa | Perturb σ=0.20, T=0.95 | 1.000 | 1.169 / 1.169 | 0.569 / 0.569 |
| E. coli vs. K. pneumoniae | Prior σ=0.30, T=0.95 | 1.000 | 2.074 / 2.074 | 0.773 / 0.773 |
| E. coli vs. K. pneumoniae | Perturb σ=0.05, T=0.75 | 1.000 | 1.169 / 1.169 | 0.563 / 0.563 |

#### SM7. SWF1 Training Protocol

The ensemble models were utilized to perform what we called surrogate-weighted fine-tuning (SWF). The balancing training dataset utilized was constructed by filtering 1,716 baseline-generated candidates through hard physicochemical constraints followed by selection of the top 30% by penalized conservative UCB score. Per-sequence training weights were derived from the same penalized score and rescaled to [0.1, 1.0]. The following hyperparameters were used:

***Table SM7. SWF1 training hyperparameters.***

| Parameter | Value |
|---|---|
| Maximum epochs | 120 |
| Early stopping | Validation loss, patience 15 |
| Best checkpoint | Epoch 6, val_loss = 3.464 |
| Learning rate | $5 \times 10^{-5}$ |
| Weight decay | 0.01 |
| Batch size | 16 |
| β (KL weight) | 0.1 |
| Free-bits | 0.10 |
| w_lm | 1.0 |
| w_kl | 0.1 |
| w_reg | 0.5 |
| Optimizer | AdamW with cosine LR decay |
| Precision | bf16-mixed |
| Val fraction | 0.10 |
| Training pool size | 478 sequences (470 unique) |
| Selection criterion | Top 30% by penalized conservative UCB score |
| Weight range | [0.1, 1.0], from penalized score |

#### SM8. CEM and RL Hyperparameter Configuration

The Cross-Entropy Method (CEM) and REINFORCE optimizers were applied with the following configurations. All CEM runs used prior-centered initialization ($\mu = 0$) unless otherwise noted.

***Table SM8a. CEM optimizer hyperparameters.***

| Parameter | Value |
|---|---|
| Iterations | 20 |
| Population size | 200 sequences per iteration |
| Elite fraction | 0.20 (40 sequences retained) |
| Elite buffer size | 500 sequences maximum |
| Smoothing factor $\alpha$ | 0.50 |
| $\mu$ initialization | 0 (prior-centered) |
| $\sigma$ initial | 0.95 |
| $\sigma$ final | 1.20 |
| Gate scale initial | 0.82 |
| Gate scale final | 1.05 |
| $\sigma$ mixing rate | 0.20 per iteration |
| Min Jaccard diversity | 0.20 (between elite sequences) |
| Temperature | 0.75 |
| top-p | 0.95 |
| Score mode | composite_80_20 (80% QSAR + 20% regression head) |
| $\kappa$ (UCB penalty) | 1.0 |

***Table SM8b. REINFORCE optimizer hyperparameters.***

| Parameter | Value |
|---|---|
| Algorithm | REINFORCE |
| Steps | 60 |
| Batch size | 8 |
| Learning rate | $2 \times 10^{-6}$ |
| KL penalty β | 0.20 |
| Optimizer | AdamW |
| Gradient clipping | 1.0 (unit norm) |
| Reward | conservative_UCB + 0.5·valid_bonus − 0.2·rep_penalty |
| Baseline decay | 0.95 (exponential running mean) |
| Reference model | Frozen copy of initialized weights |
| Elite anchors | Sampled from CEM elite buffer |
| Temperature | 0.70 |
| top-p | 0.90 |

### Supplementary Results

#### SR1. Complete Pipeline Branch Comparison

The next table correspond to a summary of different evaluated branches of the pipeline, evaluated through fixed scoring protocol coming from top 250 candidates. Besides ensemble scorer (mean score), different physicochemical parameters were extracted to provide information about the chemical behavior of the generator.

***Table SR1. Complete pipeline branch comparison (top 250 candidates, external QSAR scorer). Lower mean score indicates better predicted activity.***

| Branch | Mean score | Best | Novelty | Length | Cat. ratio | GRAVY | Interpretation |
|---|---|---|---|---|---|---|---|
| SWF1 no-CEM (prior) | 1.2767 | 1.1529 | 0.596 | 16.5 | 0.425 | — | Best prior-mode result |
| SWF1+CEM | 1.3446 | 1.1614 | 0.605 | 16.7 | 0.408 | −0.456 | BEST — recommended |
| SWF1+CEM+RL | 1.4422 | 1.4029 | 0.662 | 16.8 | 0.390 | −0.398 | Negative control |
| Baseline no-CEM (perturb) | 1.4628 | 1.3825 | 0.695 | 18.5 | 0.429 | −0.415 | Baseline reference |
| SWF1 no-CEM (perturb) | 1.4655 | 1.4189 | 0.693 | 14.9 | 0.378 | −0.525 | SWF1 without CEM |
| Baseline+CEM → SWF no-CEM | 1.5273 | 1.4500 | 0.586 | 11.7 | 0.404 | −0.965 | Pool quality insufficient |
| SWF2+CEM | 1.6849 | 1.1859 | 0.626 | 15.8 | 0.387 | −0.403 | Partial recovery only |
| Baseline+CEM | 2.3659 | 1.3222 | 0.521 | 12.5 | 0.427 | −0.589 | CEM degrades baseline |
| Baseline+RL (baseline anchors)+CEM | 2.9933 | 1.4812 | 0.503 | 12.7 | 0.412 | −0.473 | RL does not compensate |
| Baseline no-CEM (prior) | 3.0439 | 2.9164 | 0.529 | 10.9 | 0.471 | −1.098 | Poor prior baseline |
| Baseline+CEM → SWF+CEM | 3.0689 | 1.2766 | 0.522 | 13.3 | 0.424 | −0.410 | Geometry not recovered |
| SWF_prior_real+CEM | 3.1706 | 1.4910 | — | — | — | — | Not rescued by CEM |
| Baseline+RL (SWF1 anchors)+CEM | 3.2416 | 2.9626 | 0.530 | 12.9 | 0.408 | −0.483 | Anchors cannot compensate |
| SWF_prior_real no-CEM | 3.2515 | 3.1752 | — | — | — | — | Worse than baseline |
| SWF2 no-CEM | 3.5186 | 3.3269 | 0.881 | 14.8 | 0.165 | +0.138 | Catastrophic degradation |

#### SR2. CEM Convergence Data

The next table provides external evaluation for each Cross Entropy iteration regarding to the best performing model (perturb-trained SWF generation through CEM) and baseline CEM (operating directly to the decoder). QSAR mean row corresponds to the mean external score of the 40 retained elite sequences at each iteration while population mean (qm) is the mean external score across all valid sequences in the population of 200.

***Table SR2. CEM convergence: elite quality and population quality per iteration.***

| Iteration | Baseline — elite QSAR mean | Baseline — pop. mean (qm) | SWF1 — elite QSAR mean | SWF1 — pop. mean (qm) |
|---|---|---|---|---|
| 1 | 3.318 | 3.373 | 1.314 | 1.783 |
| 2 | 1.486 | 2.222 | 1.278 | 1.650 |
| 3 | 3.091 | 3.142 | 3.115 | 3.129 |
| 4 | 3.074 | 3.113 | 1.287 | 1.474 |
| 5 | 3.087 | 3.123 | 1.343 | 1.503 |
| 6 | 3.027 | 3.078 | 1.261 | 1.518 |
| 7 | 3.089 | 3.119 | 1.246 | 1.432 |
| 8 | 3.090 | 3.126 | 1.301 | 1.384 |
| 9 | 3.121 | 3.149 | 1.292 | 1.378 |
| 10 | 3.087 | 3.140 | 1.281 | 1.368 |
| 11 | 3.072 | 3.113 | 1.300 | 1.362 |
| 12 | 3.097 | 3.119 | 1.275 | 1.345 |

#### SR3. SWF2 Physicochemical Collapse

Table SR3 compares key physicochemical descriptors of generated sequences across the baseline CVAE, SWF1, and SWF2 pipelines. It illustrates how catastrophic degrading looks after the second iterative cycle of surrogate weighted fine-tuning, which decrease cationic ratio and GRAVY while it increases novelty.

***Table SR3. Physicochemical profile comparison: Baseline, SWF1, and SWF2 (perturb mode, top 250 candidates).***

| Branch | Mean length | Cat. ratio | GRAVY | Rep3 | Novelty |
|---|---|---|---|---|---|
| Baseline no-CEM (perturb) | 18.52 | 0.429 | −0.415 | 2.108 | 0.695 |
| SWF1 no-CEM (perturb) | 14.85 | 0.378 | −0.525 | 1.544 | 0.693 |
| SWF2 no-CEM (perturb) | 14.78 | 0.165 | +0.138 | 1.523 | 0.881 |

#### SR4. Top 100 SWF1+CEM Candidate Sequences

Table SR4 corresponds to the top 10 candidates generated by the best performing model (perturb-trained SWF generating through CEM). All candidates were generated against *E. coli*. A list of physicochemical descriptors was calculated utilizing modlAMP 4.3.2 to compare their chemical profile, and their pLDDT values are observable based on their ESMFold predicted three-dimensional structure. Helicity is the fraction of residues with backbone dihedral angles in the α-helical region of the Ramachandran plot ($\varphi \in [-80°, -40°]$, $\psi \in [-60°, -20°]$).

***Table SR4. Top 10 SWF1+CEM candidates by external QSAR score, with full-set summary statistics.***

| Sequence | QSAR score | Length | Net charge | GRAVY | Cat. ratio | Hyd. moment | Helicity | pLDDT |
|---|---|---|---|---|---|---|---|---|
| KWKLFKKFKLFKLKLLKLL | 1.161 | 19 | 7.99 | 0.15 | 0.421 | 0.327 | 1.00 | 90.7 |
| KKIWQKIKKFLKFLKKIF | 1.181 | 18 | 7.99 | −0.34 | 0.444 | 0.791 | 1.00 | 91.2 |
| LKWKKWLKFKWKKWLK | 1.190 | 17 | 6.99 | −0.41 | 0.412 | 0.752 | 1.00 | 88.4 |
| WRKYWKILKFL | 1.195 | 11 | 4.00 | −0.41 | 0.364 | 0.695 | 1.00 | 85.3 |
| WRKFWKYLKKFLKFL | 1.196 | 15 | 5.99 | −0.47 | 0.400 | 0.812 | 1.00 | 89.1 |
| KWKLFKKFKLFK | 1.198 | 12 | 5.99 | 0.17 | 0.417 | 0.398 | 1.00 | 87.6 |
| KFWKKWKKFLKFL | 1.199 | 13 | 5.99 | −0.23 | 0.462 | 0.721 | 1.00 | 86.9 |
| KWKIFRWWKFLK | 1.204 | 12 | 5.00 | −0.17 | 0.417 | 0.698 | 0.83 | 82.1 |
| KRIVQRIKKFLKFL | 1.205 | 14 | 7.00 | −0.29 | 0.500 | 0.683 | 1.00 | 88.3 |
| LKWWLKWLKFKK | 1.208 | 12 | 5.00 | −0.25 | 0.333 | 0.745 | 1.00 | 84.7 |
| …(90 additional sequences) | 1.21–1.43 | 8–25 | 3–9 | varies | 0.15–0.55 | 0.3–0.9 | 0–1.0 | 65–95 |

Summary statistics for the full set of 100 candidates: mean QSAR score 1.259, best 1.161, top-10 mean 1.195, mean length 17.2, mean net charge 6.83, mean GRAVY −0.421, mean cat. ratio 0.416, mean hydrophobic moment 0.598, mean helicity 0.874, 96% of candidates with helicity > 0.5, mean pLDDT 83.7, ESMFold success rate 100%.

#### SR5. Independent APEX Validation: Full Results

Table SR5 reports the mean predicted activity (μg/mL) of four of our models by APEX, to compare the best-performing model against the perturb, prior and CEM baselines top 100 candidates. APEX is a multitask deep learning model from the Machine Biology Group of the University of Pennsylvania. Spearman correlations between APEX and our models were not found (were r = 0.06, 0.21, and −0.09 for E. coli, K. pneumoniae, and P. aeruginosa respectively), confirming linear independence between the two scoring systems

***Table SR5. APEX model predicted MIC (μg/mL) across four pipeline configurations and three target organisms. Lower values indicate better predicted antimicrobial activity.***

| Organism | SWF1+CEM (best) | Baseline quality-first | Baseline balanced | Baseline prior (exploration) |
|---|---|---|---|---|
| E. coli | 34.4 | 44.4 | 37.1 | 54.6 |
| K. pneumoniae | 132.9 | 184.6 | 186.2 | 176.8 |
| P. aeruginosa | 59.5 | 74.9 | 71.0 | 87.1 |
